# Possible role of left–right asymmetry in the sensory system and behavior during adaptation to food-sparse cave environments

**DOI:** 10.1101/2022.07.09.499441

**Authors:** Vânia Filipa Lima Fernandes, Yannik Glaser, Motoko Iwashita, Masato Yoshizawa

**Affiliations:** School of Life Sciences, University of Hawai‘i at Mānoa, Honolulu, Hawai‘i, USA; Department of Information and Computer Sciences, University of Hawai‘i at Mānoa, Honolulu, Hawai‘i, USA

## Abstract

Laterality in relation to behavior and sensory systems is found commonly in a variety of animal taxa. Despite the advantages conferred by laterality (e.g., the startle response and complex motor activities), little is known about the evolution of laterality and its plasticity in response to ecological demands. In the present study, a comparative study model, the Mexican tetra (*Astyanax mexicanus*), composed of two morphotypes, i.e., riverine surface fish and caved-welling cavefish, was used to address the relationship between environment and laterality. The use of a machine learning-based fish posture detection system and sensory ablation revealed that the left cranial lateral line significantly supports one type of foraging behavior, i.e., vibration attraction behavior, in one cave population. Additionally, left–right asymmetric approaches toward a vibrating rod became symmetrical after fasting in one cave population but not in the other populations. Based on these findings, we propose a model explaining how the observed sensory laterality and behavioral shift could help adaptations in terms of the tradeoff in energy gain and loss during foraging according to differences in food availability among caves.

## Introduction

Laterality, which is common among various animal taxa, is expressed, for example, as the asymmetrical use of the left or right hand or foot (Tommasi, 2009), the biased use of the left or right eye, and the use of the left or right brain hemispheres for different processing tasks (Heilman and Abell, 1980; Nottebohm, 1977, 1971; Nottebohm and Nottebohm, 1976; Rogers, 1981; Rogers and Workman, 1989; Schwartz et al., 1975; Wernicke, 1874). Laterality confers several advantages in recognizing and escaping from predators (Chivers et al., 2017, 2016; Dadda et al., 2010), facilitating complex and intricate motor activities (Magat and Brown, 2009) and enabling spatial learning (Sovrano et al., 2005). Phylogenetic studies have shown that left–right (L–R) asymmetric traits have repeatedly and independently evolved among different animal taxa (Güntürkün et al., 2020), possibly due to the need for laterality in diverse environments. However, the influence of environments on variation in the laterality and its plasticity among populations has not been well-studied.

As an entry point to solve this evolutionary question, we explored the environment–laterality relationship in *Astyanax mexicanus*, the Mexican tetra, which has two interfertile morphotypes, i.e., surface fish and cavefish (Keene et al., 2016; Protas and Jeffery, 2012; Rétaux et al., 2013; Wilkens et al., 1988) and is known as a model for studying the evolution of behavior. Cavefish, the blind cave-dwelling form of *A. mexicanus*, were separated from their surface-dwelling ancestors about 2,000–20,000 years ago (Fumey et al., 2018; Herman et al., 2018). In total, 32 *A. mexicanus* cave populations have been described in Mexico (Espinasa et al., 2020), and many of these populations have evolved morphological, physiological, and behavioral traits independently, likely via adaptation (Borowsky, 2008; Mitchell et al., 1977; Ornelas-García et al., 2008; Strecker et al., 2012; Wilkens et al., 1988).

One type of laterality has been reported in the Pachón cave population of *A. mexicanus* cavefish, i.e., a preference for using the right-side of the body to detect novel immotile objects in the environment (de Perera et al., 2005). However, it is not yet known which environmental factors determine this type of laterality (unclear ecological relevance) or whether it evolved in a surface fish-like ancestor because such laterality in surface fishes has not been investigated. In the present study, we investigated a well-characterized type of foraging behavior, vibration attraction behavior (VAB), which is evoked in response to 35–40 Hz vibrations (Yoshizawa et al., 2010) within the range potentially produced by soil crustaceans and percolating water (Lang, 1980; Montgomery and Macdonald, 1987). VAB is evolutionarily enhanced in cavefish, and its advantage was demonstrated as they hunt small prey in the dark (Yoshizawa et al., 2010). Different degrees of increased VAB levels have been reported in lab cave populations and in the original wild cave populations (Espinasa et al., 2021; Yoshizawa et al., 2010), likely due to differences in ecological factors in these caves. Outside of cave environments, VAB may be detrimental because of nocturnal predators, such as prawn fish (Wilson et al., 2004; Yoshizawa and Jeffery, 2011).

VAB is mediated by the mechanosensory lateral line system, which is composed of sensory units known as superficial neuromasts (SNs) particularly located at the eye orbit and the third infraorbital bone (IO3) (Yoshizawa et al., 2012b, 2010). We reported previously that, despite L–R symmetry in the number of SNs in surface and cave morphs, a positive correlation existed between the number of left SNs and VAB level in Pachón cavefish (Fernandes et al., 2018). This asymmetric sensory usage (hereafter termed “sensory laterality”) was not observed in surface fish, suggesting that this sensory laterality emerged through an evolutionary process that conferred some advantage to fish that colonized cave environments (Fernandes et al., 2018).

In the current study, we investigated the asymmetric L–R responses to a vibrating stimulus in *A. mexicanus* populations. In contrast to the Pachón cavefish, independent cavefish populations from the Los Sabinos and Tinaja caves did not show detectable coupling between the left or right SN and VAB levels, indicating that asymmetric sensory–behavior coupling could be adaptive in the Pachón cave. SN ablation revealed that the left SN in Pachón cavefish indeed regulate the number of approaches (NOA) level variation. In Tinaja cavefish, we found alternative L–R asymmetry in their approaches toward a vibrating rod that did not depend on the number of L–R SNs but was dependent on a fasting condition. Specifically, Tinaja individuals that exhibited asymmetric L–R approaches prior to fasting showed L–R balanced approaches after fasting. Such fasting-dependent L–R plasticity was not detected in the Pachón cavefish or surface fish populations.

Our results indicate that different cave populations of *A. mexicanus* have evolved distinct laterality strategies. Our study also suggests that L–R sensory usage and the L–R asymmetric approach have different underlying mechanisms. Based on these findings, we propose a model that explains the advantages of laterality according to diet availability and energy gain and loss. This model implies that the laterality of Pachón cavefish saves energy most effectively among the studied cave and surface populations and may have allowed Pachón cavefish to adapt successfully to the fluctuating availability of food in the Pachón cave.

## Results

### Sensory laterality among different cave populations of *A. mexicanus*

In a previous study, we reported a positive correlation between the VAB level and the number of left- but not right-side SNs in Pachón cavefish (Fernandes et al., 2018). This L–R asymmetric association between the number of SNs and VAB likely evolved during the cave adaptation process. However, whether such left SN–VAB coupling exists in other discrete cave populations in the context of convergent/parallel evolution was unclear. Thus, we investigated other VAB-positive cave populations in Tinaja and Los Sabinos (Espinasa et al., 2021; Yoshizawa et al., 2015, 2010) (Figure 1). First, to detect left or right approaches toward a vibrating rod, we developed a method using DeepLabCut (DLC), a machine-learning suite (Mathis et al., 2018; see Materials and Methods), with which we accurately measured the X–Y coordinates of the left and right craniums of fish, which was not possible with a method we developed previously (Yoshizawa et al., 2015, 2010). To validate the new method, we measured the number of approaches toward the vibrating rod (NOA) and the duration around the vibrating rod (DIR) in one surface fish and three cavefish populations (see Materials and Methods). Consistent with previous results (Yoshizawa et al., 2015, 2012a, 2010), Pachón cavefish showed the highest level of VAB in comparison to Los Sabinos and Tinaja cavefish and surface fish, both in terms of NOA and DIR (Figure 1C; Supplementary Table 1), indicating that the new detection method is consistent with the method developed previously (Yoshizawa, 2015; Yoshizawa et al., 2012a, 2010).

**Figure 1.**
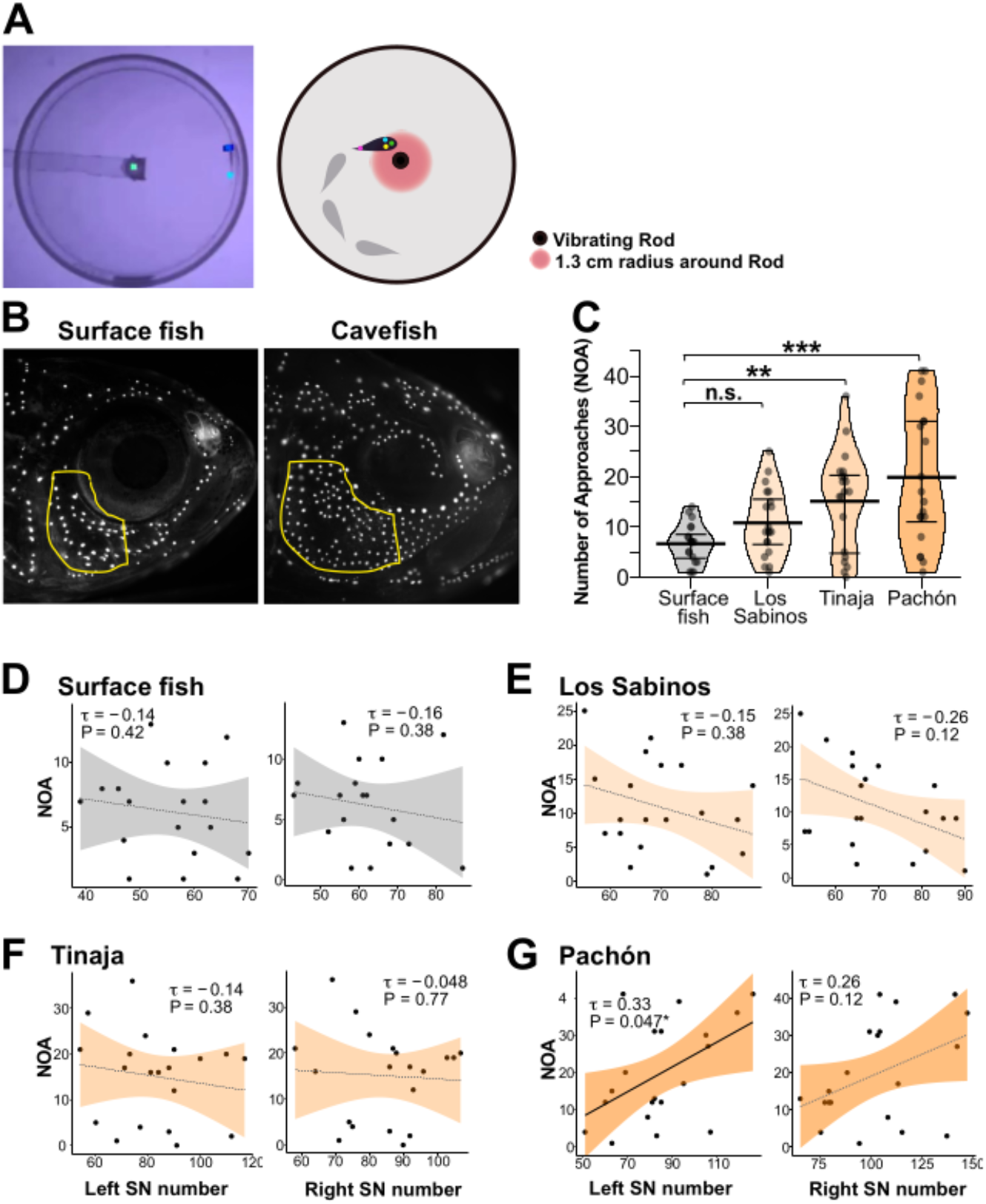
Sensory laterality in surface fish and cavefish populations of *Astyanax mexicanus*. VAB videos were analyzed using a home-developed Python script and a DeepLabCut deep learning algorithm. This method allowed us to detect the number of times fish made left or right approaches toward the vibrating rod (NOA) and the associated duration they spent within a 1.3-cm radius from the vibrating rod (DIR). (**A**) Video image (left) and schematic representation (right) of fish and the vibrating rod labeled with three markers on the head (left, center, and right) and one marker on the caudal fin. (**B**) Superficial neuromasts (SNs) in the infraorbital region (IO3 region delimited by a yellow line) were stained with 4-Di-1-ASP vital dye. (**C**) NOA data for the lab-raised populations indicated that Pachón cavefish showed the highest NOA values. The bars in the pirate plots represent means ± standard errors of the means (each population, n = 20). *, P < 0.05; **, P < 0.01; and ***, P < 0.001. Statistical scores are shown in Supplementary Table 1. (**D–G**) The number of SNs in the left or right IO3 area plotted against the total NOA in surface fish (**D**) and Los Sabinos (**E**), Tinaja (**F**), and Pachón cavefish (**G**). Statistical scores are available in Supplementary Table 2.

We then explored whether an L–R bias existed in the couplings between the number of SNs and NOA or DIR. In the surface population, no correlation was detected between the left SNs and the L–R summed NOA/DIR (i.e., total NOA/DIR) or the right SNs and the total NOA/DIR (Figure 1D; statistics in Supplementary Table 2; all NOA and DIR in Figs. 1 and 2 are L–R summed). In contrast, a positive correlation was found between the left SNs and NOA in the Pachón cave population (τ = 0.329, P = 0.047; Figure 1G; Supplementary Table 2) but not between the right SNs and NOA (τ = 0.258, P = 0.118; Figure 1G; Supplementary Table 2). This result is also consistent with our previous study (Fernandes et al., 2018); thus, we tested the other cave populations with confidence. Both the Los Sabinos and Tinaja cave populations showed results similar to those of the surface population, i.e., no correlation was detected between the left SNs and NOA/ DIR or between the right SNs and NOA/ DIR (Figure 1E and F; Supplementary Table 2). In summary, our machine-learning-based detection method generated results consistent with our previous studies, showing that the left SN number is coupled with the NOA level in the Pachón cave population. However, this method failed to detect such correlations in other cave populations.

**Figure 2.**
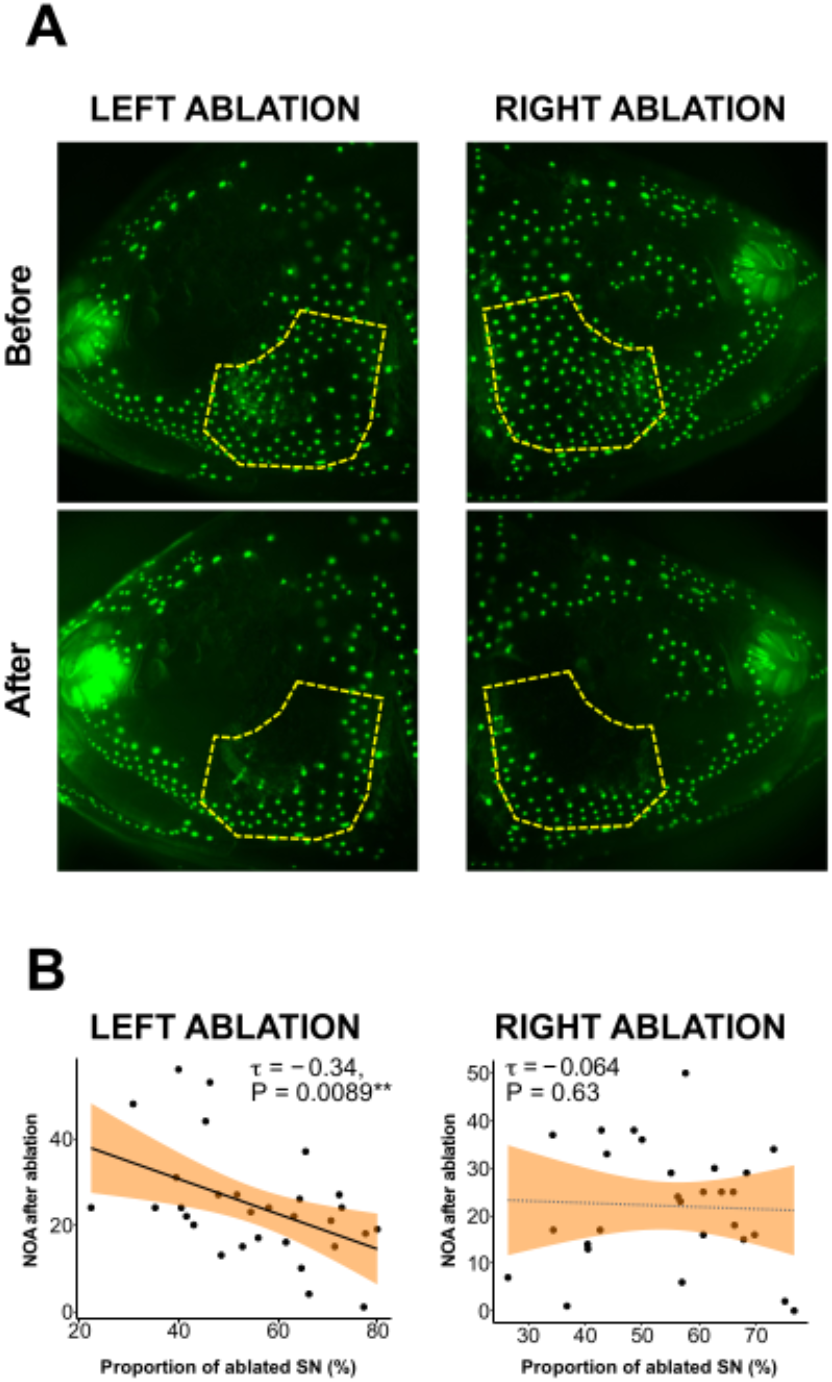
Ablation of superficial neuromasts (SNs) confirmed that the left-side SN promotes the level of VAB in Pachón cavefish. (**A**) Left and right sides of SNs in Pachón cavefish stained with 4-Di-1-ASP (green dots) before and after ablation of SNs in the infraorbital region (IO3 shown by a yellow line) of the same fish. (**B**) Proportion of ablated SNs on the left-side and right-side plotted against the total NOA in both cases. Regression lines with 95% confidence intervals (shaded gray or orange) are shown in each panel. See Supplementary Table 3 for additional details. **, P < 0.01.

### Coupling between the left SNs and VAB is based on inputs to the left SNs

To investigate the underlying mechanism of the coupling between the left SNs and VAB, we ablated the SNs on either the left or right-side at the IO3 area in the Pachón cave populations using a well-developed noninvasive method (Yoshizawa et al., 2010). To highlight SN–VAB coupling, we selected VAB-positive individuals [NOA ≥ 6; (Yoshizawa et al., 2010)]. The ablation was confirmed using DASPEI staining after the behavior assay (Figure 2A and B). After ablation, Pachón cavefish individuals with a higher percentage of ablated SNs exhibited lower levels of VAB (τ = −0.342, P = 0.009; Figure 2B; Supplementary Table 3). However, the change in VAB was not observed when the right-side SNs were ablated (τ = −0.064, P = 0.635; Figure 2D; Supplementary Table 3). We also performed SN ablation in surface fish and detected no correlation between NOA and the proportion of ablated SNs on the left or right sides (Figure 2—figure supplement 1; Supplementary Table 3). These results reinforce our previous findings (Fernandes et al., 2018) regarding the significant role of the left-side SNs in the VAB of the Pachón cave population.

For DIR (i.e., the time spent located within 1.3 cm of the vibrating rod), no correlation was found with the proportions of left or right ablated SNs in Pachón cavefish (Supplementary Table 3), suggesting a possibility that the NOA and DIR are regulated differently, i.e., in Pachón cavefish, NOA could be mediated by the SNs, whereas DIR could be regulated via non-SN means, such as auditory and/or tactile sensing (see the Discussion).

Regarding population averages, for Pachón cavefish, the ablation of the left, right, or left and right SNs induced no detectable changes in NOA or DIR (although the ablation of left-side SNs increased NOA significantly: V = 74.5, P = 0.032; Figure 2—figure supplement 2; Supplementary Table 4; See also Supplementary file 1—extended result and discussion). In summary, for Pachón cavefish, the left SN number raises NOA but not DIR, and the right SNs were not significantly associated with NOA.

### Behavioral laterality across different populations of *A. mexicanus*

Our results showed that the left SNs promoted VAB in Pachón cavefish, whereas the other populations did not show L–R-biased coupling between the SNs and VAB (i.e., sensory laterality) (Figs. 1 and 2). Given that VAB evolved as a foraging behavior and that diet type and availability differ among caves [e.g., from decomposed plants to soil or cave-adapted arthropods as well as seasonal flooding (Culver, 1982; Culver and Pipan, 2009; Espinasa et al., 2017; Keene et al., 2016; Mitchell et al., 1977; Yoshizawa et al., 2010; Yoshizawa and Jeffery, 2011)], we hypothesized that alternative forms of laterality in VAB exist in surface and/or cave populations. Thus, we explored two types of L–R bias in VAB (i.e., left/right NOA and left/right DIR; Figure 3A) among the surface population and Pachón, Tinaja, and Los Sabinos cave populations. To quantify the lateral preference of individual fish, the right NOA or DIR was divided by the total NOA or total DIR, respectively (i.e., ratio scores of >0.67 and <0.33 indicate right and left preferences, respectively; Figure 3B and 3C). We measured the L–R biases of NOA and DIR separately because our results indicated that they were likely regulated differently (see the previous section and Discussion). Individuals from each of the cave and surface populations showed a preference for one side or the other while approaching (NOA) and touching the rod (DIR) (Figure 3B and 3C, respectively). This L–R preference in each individual was not repeatable or only fairly repeatable in surface fish and Pachón cavefish when we conducted assays 6 days apart and repeated three times (Table 1) [respective NOA intraclass correlation coefficients (ICCs): κ = 0.0 (none), P = 0.48; κ = 0.39 (fair), P = 0.00025. Respective DIR ICCs: κ = 0.0 (none), P = 0.48; κ = 0.085 (none), P = 0.20]. In contrast, Tinaja cavefish individuals exhibited a moderate level of repeatability in NOA (κ = 0.54, P = 0.0021; repeated three times; Table 1), suggesting that these fish tended to consistently approach the vibrating rod unidirectionally.

**Figure 3.**
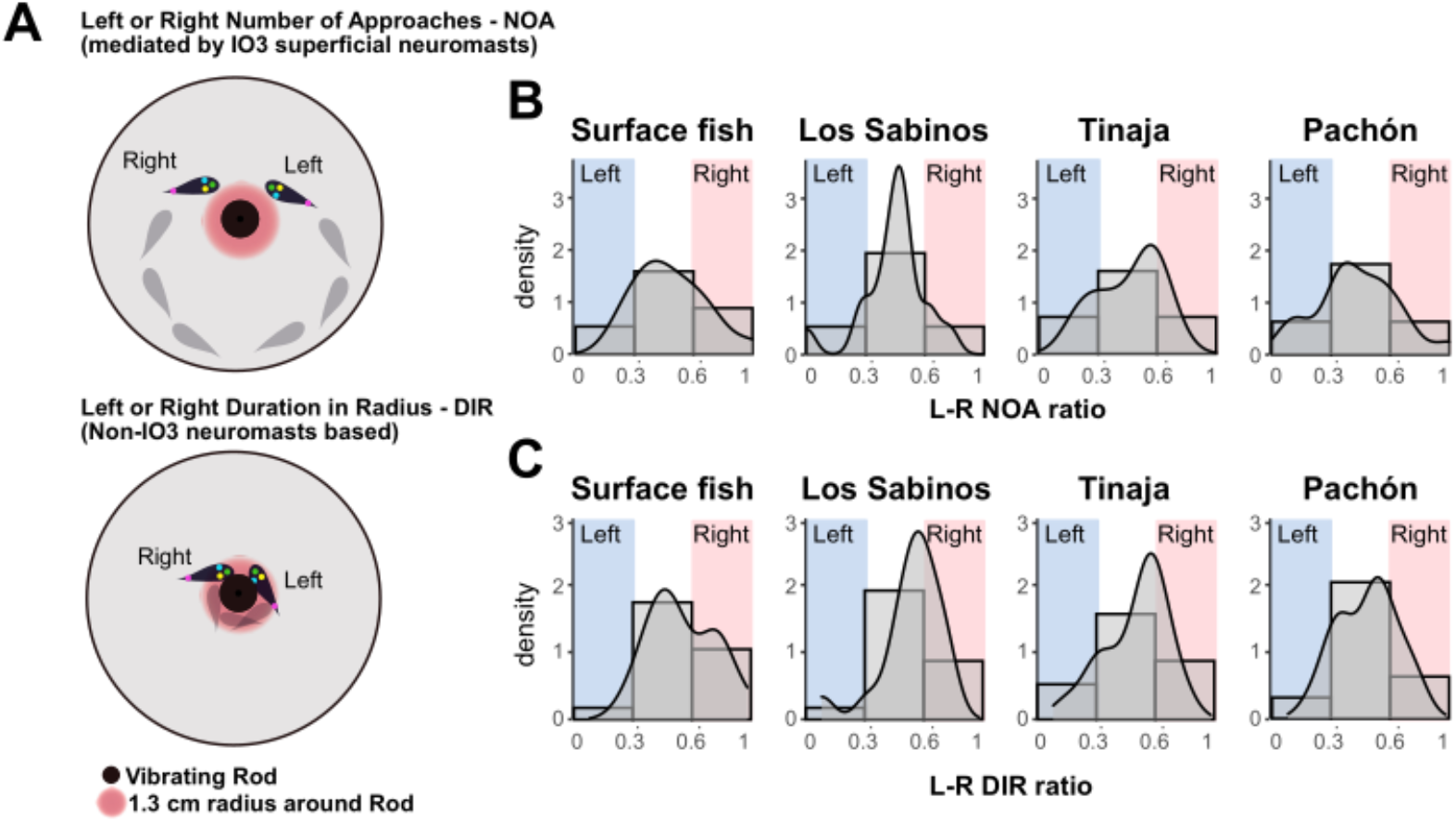
Laterality in behavioral output among surface fish and cavefish populations of *Astyanax mexicanus*. (**A**) Schemes representing left or right approaches expressed as NOA (top) and left or right adherence expressed as DIR (bottom). (**B**) Histograms for the L–R NOA ratio (the right-side NOA divided by the total 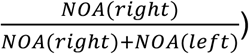 across all four populations. (**C**) Histograms for the L–R DIR ratio 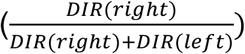 across all four populations. Ratios of <0.33 indicate a strong left preference, ratios of 0.33–0.67 indicate balanced approaches, and ratios of >0.67 indicate a strong right preference. Los Sabinos cavefish showed a gaussian-like distribution, whereas many surface fish and the other cavefish individulas exhibited strong left or right bias. Notably, all four populations included a few individuals that showed strong left or right preferences (near 0.0 or 1.0, respectively).

**Table 1.** Intraclass correlation coefficient (ICC) under Cohen’s suggestion (Landis and Koch, 1977; Zegers et al., 2010) (N = 3 repeats; N= 22 for surface, N = 9 for Pachón cave and N = 11 for Tinaja cave fish).

| Population | Measurement | Average L–R ratio | Statistics | Kappa | P-values |
| --- | --- | --- | --- | --- | --- |
| Surface fish | NOA | $0.459 \pm 0.034$ | $F_{21,42} = 1.0$ | 0.0 (none) | 0.48 |
| Surface fish | DIR | $0.481 \pm 0.028$ | $F_{21,42} = 1.0$ | 0.0 (none) | 0.48 |
| Pachón cavefish | NOA | $0.464 \pm 0.036$ | $F_{28,56} = 3.0$ | 0.39 (fair) | 0.00025*** |
| Pachón cavefish | DIR | $0.473 \pm 0.024$ | $F_{28,56} = 1.3$ | 0.085 (none) | 0.20 |
| Tinaja cavefish | NOA | $0.482 \pm 0.052$ | $F_{10,20} = 4.5$ | 0.54 (moderate) | 0.0021** |
| Tinaja cavefish | DIR | $0.478 \pm 0.046$ | $F_{10,20} = 2.4$ | 0.32 (fair) | 0.043* |

As shown in the histograms in Figure 3B and C, there was no detectable bias between the left and right-side NOA (F_1,152_ = 0.149, P = 0.700) or between the left and right DIR among the populations (F_1,152_ = 3.378, P = 0.068) (Figure 3—figure supplement 1; Supplementary Table 1). Additionally, there was no detectable bias between the duration of left and right facing outside the 1.3-cm radius around the vibrating rod, suggesting that there was no tendency to face left or right in relation to the vibrating rod from a distance (F_3,152_ = 0.016, P = 0.997; Supplementary Table 1).

The above results suggested that laterality in SN usage (sensory: Pachón cavefish) and laterality in approaches (behavior: Tinaja cavefish) existed in some *A. mexicanus* populations. Therefore, we investigated whether relationships existed between the lateralities in the number of left or right SNs and left or right NOA or left or right DIR. Detailed correlation analyses were performed in the ablation study. In Pachón cavefish, the ablation ratio of the left SNs mirrored the reduction of left approaches (left NOA), but no correlation was detected between the left SN ablation ratio and the right NOA, the right SN ablation ratio and the left or right NOA, and the left or right SN ablation ratio and the left or right DIR (Supplementary Table 3). Additionally, there was no detectable effect of any left or right SN ablation on the L–R approaches (NOA or DIR) of surface fish. These results indicate that the number of left SNs (inputs) affects the left approaches of Pachón cavefish but not those of surface fish. As in the intact SN condition, there were some positive correlations between the number of L–R SNs and the L–R NOA and/or DIR in surface fish and Pachón cavefish (but not in other cave populations; see Supplementary Table 2). However, the SN–behavior relationship of surface fish in the intact state may not be vital because it was not detected in the ablation study.

Collectively, these results indicate that sensory laterality was found only in Pachón cavefish and was associated with left-side approaches. Individuals in the Tinaja cave population but not in the surface or Pachón cave populations showed a moderately consistent preference for left- or right-side approaches, and L–R bias was not detectable at the whole population level.

### Fasting increased sensory responses and reduced approach laterality

Previous prey-capture experiments indicated that VAB promotes the food capture rate in dark and food-sparse environments (Yoshizawa et al., 2010). VAB plasticity according to the environment has also been suggested (Espinasa et al., 2021). Thus, we hypothesized that fasting induces shifts in laterality. Because younger fish are more sensitive to fasting in general, we used 1-year-old (young adult) surface fish, Pachón cavefish, and Tinaja cavefish rather than the 2–3-years-old (adult) fish used in the earlier experiments (Figs. 1–3).

First, we investigated the effect of fasting on sensory laterality (i.e., the number of left or right SNs vs. the total NOA or DIR) among the surface, Pachón, and Tinaja populations. The 6-day fasting reduced the total NOA and total DIR in surface fish but increased the total DIR in Tinaja cavefish (Figure 4A and 4B; Supplementary Table 5); however, no changes were detected in Pachón cavefish (Figure 4A and 4B; Supplementary Table 5). To investigate L–R SN–behavior coupling, we determined the correlations between either the number of left or right SNs and the total NOA after fasting with subtraction of the prefasting NOA (Figure 4C; Supplementry Table 7). Pachón individuals with fewer “left” SNs exhibited a higher increase in total NOA (Figure 4C; Supplementry Table 7). Pachón individuals with fewer “right” SNs showed a higher increase in NOA but not at a significant level (Figure 4C; Supplementry Table 7). Surface fish or Tinaja cavefish did not show fasting-dependent changes in SN–approach coupling (Figure 4C; Supplementry Table 7). In summary, fasting yielded different responses in the surface and Pachón populations, i.e., NOA was reduced in surface fish, and Pachón individuals with fewer left SNs showed a higher increase in NOA relative to that of individuals with more left SNs. Second, we studied the effect of fasting on laterality in approaches. The surface and Tinaja populations had more number of younger fishes (1-year-old fish vs. 2–3-years-old fish) that showed L–R-biased approaches in the prefasting condition (NOA; broader standard variations; Figure 5A and 5B). In contrast, Pachón cavefish individuals showed L–R symmetrical NOA with a narrower standard variation than that observed in the surface and Tinaja populations in the prefasting condition (Figure 5A and 5B). After 6 days of food deprivation, the distributions of the L–R preference in surface fish became more uniform for both NOA and DIR (Figure 5A, 5B, 5C, and 5D). Interestingly, Tinaja cavefish shifted from a right-biased approach (for both NOA and DIR) to a symmetrical approach in NOA (~0.6 in a bias index; Figure 5C) and a reduced right bias in DIR (Figure 5). Regarding population averages, none of surface fish, Tinaja, or Pachón cavefish individuals showed detectable L–R bias in NOA or DIR in the prefasting or postfasting condition, except DIR in Tinaja cavefish (right bias; Figure 5—figure supplement 1A and 1B), which mirrors the observation in Figure 5A and 5B. Fasting also induces alternative responses among surface fish, Tinaja, and Pachón cavefish individuals; surface fish decreased the overall NOA and DIR after fasting, while Tinaja cavefish increased DIR (Figure 5—figure supplement 1A and 1B). Pachón cavefish did not show detectable changes in the overall NOA and DIR levels.

**Figure 4.**
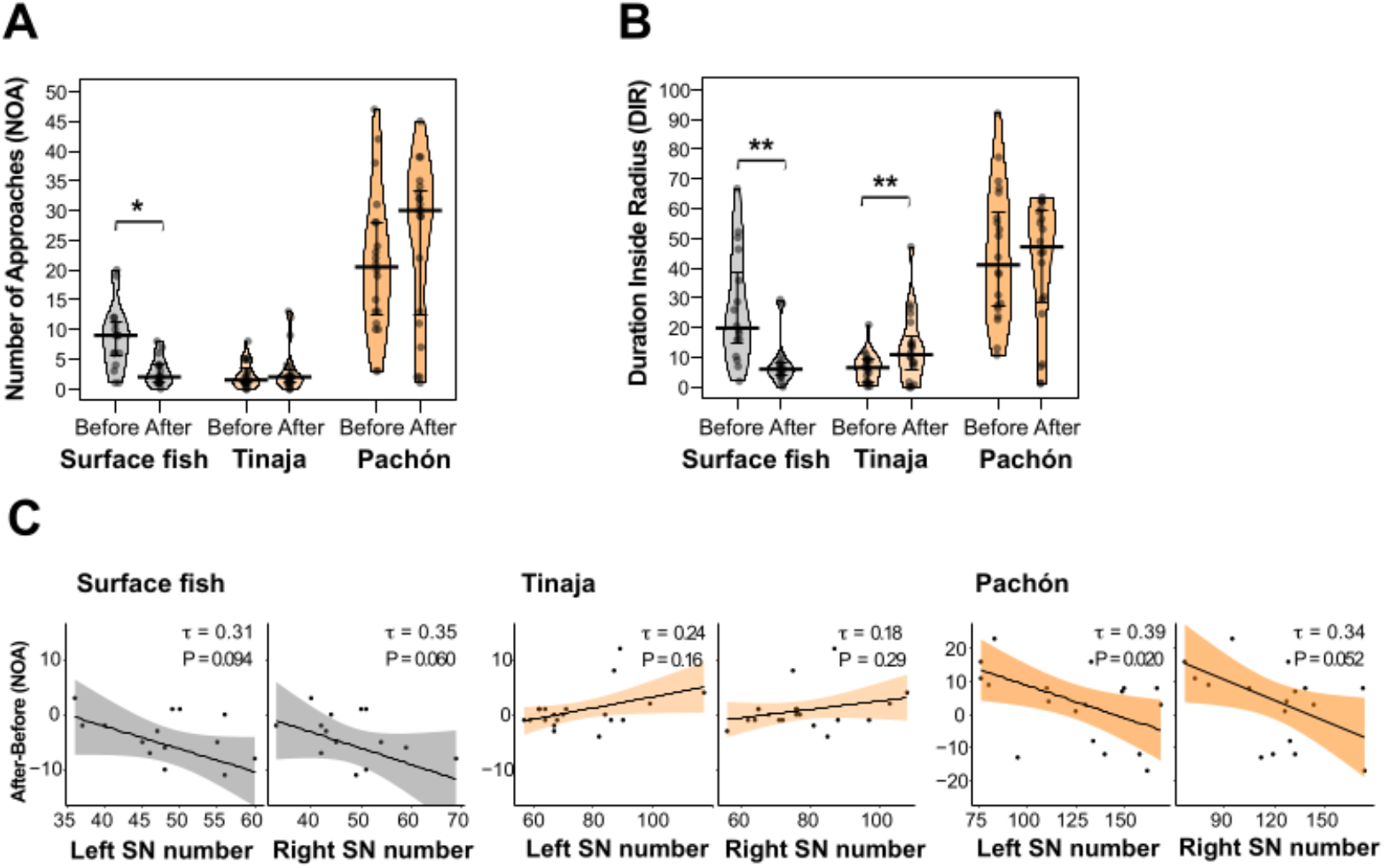
Plasticities of surface fish and Tinaja cavefish in total NOA and/or DIR, and the plastic left SN–NOA association in Pachón cavefish in response to fasting. (**A**) Pirate plots showing total NOA. Surface fish showed significantly reduced NOA after fasting, but neither Tinaja nor Pachón cavefish showed significant changes in NOA. (**B**) Pirate plots showing total DIR. Surface fish showed significantly reduced DIR after fasting, but Tinaja’s DIR increased after fasting. Pachón cavefish showed no significant changes in DIR. (**C**) The number of left or right SNs plotted against changes in NOA after fasting (NOAafter–NOAbefore) in surface fish, Tinaja cavefish, and Pachón cavefish. In Pachón cavefish, individuals with fewer left SNs showed a higher increase in total NOA, suggesting that a plastic response to fasting existed in individuals with few left SNs and a low NOA level. There was no detectable correlation between the number of right SNs and the change in NOA in Pachón cavefish or the other populations. All statistical scores are available in Supplementary Table 5 and 7.

**Figure 5.**
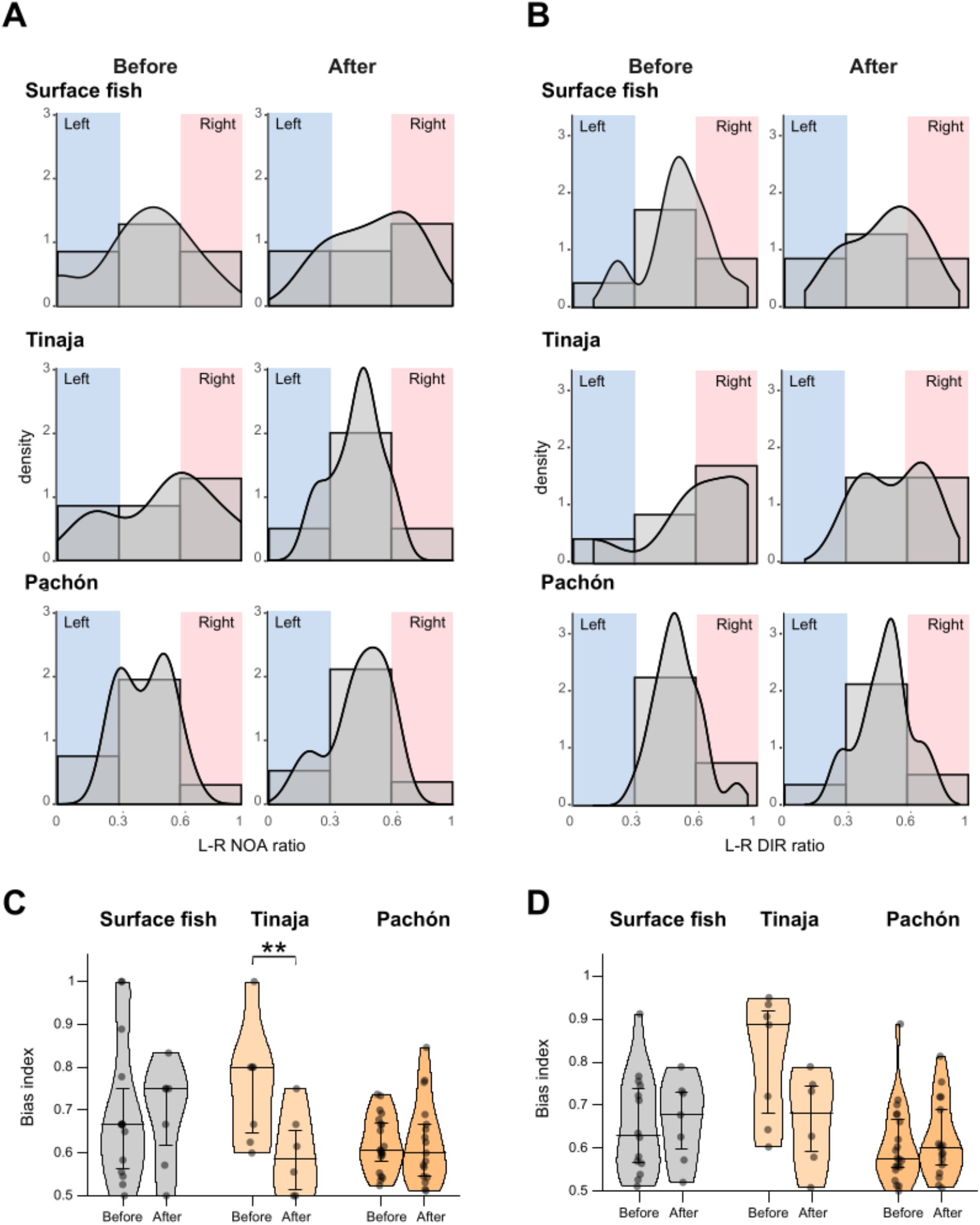
Right-biased approaches (NOA) in Tinaja cavefish were reduced after fasting. (**A, B**) Histograms and density plots showing the L–R NOA ratio (**A**) and L–R DIR ratio (**B**) before and after fasting in the indicated populations. In **A**, more Tinaja cavefish individuals showed L–R balanced approaches (0.33–0.67) after fasting; Pachón cavefish maintained L–R balanced approaches (0.33–0.67) before and after fasting, whereas surface fish showed a slight right bias. L–R NOA 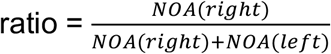. In **B**, the trends for DIR were similar to those in (A). L–R DIR ratio 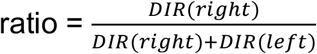. (**C, D**) Pirate plots showing L–R bias index for NOA (**C**) and DIR (**D**). In **C**, Right bias 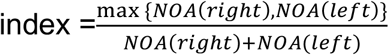. Tinaja cavefish showed a significantly reduced bias index for NOA after fasting, indicating balanced L–R approaches, whereas the other populations showed no detectable changes. In **D**, L–R bias 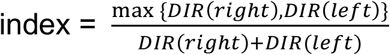. None of the populations showed detectable changes in the bias index after fasting, although Tinaja cavefish showed a nonsignificant trend for reduced bias after fasting. All statistical scores are available in Supplementary Table 6. **, P < 0.01.

In summary, food deprivation drove behavioral shifts in the surface and Tinaja cave population but not in the Pachón cave population, i.e., surface fish showed reduced VAB, whereas Tinaja cavefish showed increased VAB (Figure 5, and Figure 5—figure supplement 1). Furthermore, Tinaja cavefish showed a postfasting plastic change in the L–R bias of NOA toward a more L– R symmetric approach (Figure 5, and Figure 5—figure supplement 1).

## Discussion

We addressed laterality in sensory–behavior coupling and rod approaches among different populations of *A. mexicanus.* Using a newly developed DLC- and machine learning-based video analysis method, we found the following: (1) only the Pachón cave population showed a bias in SN–VAB coupling, i.e., the left SNs promote VAB, but not detectable in other populations; (2) no L–R bias detected in approaches (NOA) and adherence (DIR) in the surface, Pachón, Tinaja, and Los Sabinos cave populations at the population levels, but some individuals, especially those in the Tinaja cavefish population, repeatedly showed a preference for left or right approaches; and (3) fasting for 6 days reduced the L–R-biased in approaches in the Tinaja cave population, indicating that VAB plasticity is present in this population. Thus, L– R asymmetry in foraging behavior seems to have evolved flexibly under different caves, and our findings provide insights into the evolutionary dynamics of L–R asymmetry under ecological pressure (see below). At the end, we discuss the energy-saving advantage associated with foraging laterality.

### Introducing a machine-learning method to assess adaptive foraging behavior

With the aid of a DLC neural network (trained with 807 images), we developed a new video analysis method that achieved a detection accuracy of >99%, which was confirmed using a manual survey of the labeled videos. In comparison with our previously reported method (Yoshizawa et al., 2010), this trained network yielded higher resolution for the approach (NOA) and adherence (DIR) events of VAB, and left or right facing in the direction of the vibrating rod could be monitored. The new method was also confirmed to be consistent with the ImageJ-based method developed previously (Figure 1), which supports the findings of the present study as an expansion of the previous results.

### Coupling between the left SNs and NOA but not DIR in Pachón cavefish

Pachón cavefish showed a significant correlation between the number of left SNs and NOA. This coupling was observed repeatedly in different cohorts from different Pachón parents (Fernandes et al., 2018). Furthermore, this sensoory laterality in NOA in Pachón cavefish was confirmed in our ablation experiment, i.e., as more left SNs were ablated, the NOA was reduced. Notably, a correlation between the left SNs and DIR was not detected in Pachón cavefish. Such variation in sensory–behavior coupling suggests that differences exist in the sensory mechanisms underlying NOA and DIR, i.e., given the correlation between the SNs and NOA and our SN ablation results, NOA in Pachón cavefish is likely enhanced by SNs. Contrastingly, DIR was neither correlated with the number of SNs nor changed after SN ablation, indicating that other sensory systems, e.g., tactile and/or auditory sensors, affect DIR. The auditory inner ear can sense near field particle motion based on the motion lag between the fish body and the otolith (Kalmijn, 1988; Popper and Fay, 2011; Tricas, 2020). Overlapping functions between auditory, lateral line, and tactile senses have been noted but not studied in detail in the context of foraging (e.g., Braun and Sand, 2013; Kohashi et al., 2012). VAB thus emerges as a good model behavior of the modality switch between these three sensory systems because it can be evoked from the far field (>5 m in the field: auditory; see below; Espinasa et al., 2021) and near field (possibly; >1.3 cm: lateral line; <1.3 cm: tactile).

### Laterality in cranial morphology and behavior

There are reports of cranial bending in adult Pachón and Tinaja cavefish but not in adult surface fish or younger individuals of the Pachón, Tinaja, and surface populations (Powers et al., 2017). We did not detect left cranial bending in our experimental fish (2–3 cm standard length; 1–3 years old), which excludes the confounding effect of head bending on lateralities in sensory–behavior coupling and approaches to the rod. Other morphological asymmetries have been reported previously, e.g., L–R asymmetry in the suborbital (i.e., infraorbital) bone fragmentations in Pachón and Tinaja cavefish but not in surface fish (Gross et al., 2016); and L–R asymmetry in cranial SN distribution but not the number of SNs, which corresponds to asymmetric bone fragmentation (Gross et al., 2016; Gross and Powers, 2016). In the present study, we investigated the L–R asymmetric “number” but not “distribution” of SNs. It will be interesting to investigate whether the SN distribution pattern plays a role in VAB expression in future studies.

### Laterality of approaches at the population and individual levels, and the effect of starvation

In our L–R approach study, there was no detectable left or right bias for NOA and DIR at the population level among the four tested populations (with the exception of 1-year-old Tinaja cavefish). However, some individual fish showed one side bias in NOA and/or DIR, and these asymmetric biases were consistent within each individual. Laterality at the individual level has been largely overlooked (Rogers, 1989); nonetheless, studies in chicks and fish showed that behavior lateralization was random in a population but existed and seemed advantageous (e.g., by minimizing the use of brain resources) at the individual level in complex tasks, including social interactions (Bisazza et al., 2000; Bisazza and Dadda, 2005; Rogers and Workman, 1989). In the current study, we also revealed plastic laterality in a cave population responding to fasting. Because VAB evolved as a foraging behavior under food-limited conditions (Yoshizawa et al., 2010; Yoshizawa and Jeffery, 2011), we propose that cavefish laterality evolved to conserve energy.

Many caves in North-eastern Mexico experience 6-months wet and 6-months dry seasons; thus, cavefish experience food-sparse conditions for ~6 months per year (Espinasa et al., 2017). We tested the effect of food deprivation on L–R asymmetry, and removing food intake for 6 days increased both DIR and L–R symmetric NOA in the starvation resistant Tinaja cavefish (Aspiras et al., 2015). The increases in DIR and symmetric NOA could help boost food intake while saving energy (see below). In contrast, the surface fish, which are less tolerant to starvation, showed decreased NOA and DIR, seemingly saving energy in a simpler manner by reducing the number of foraging attempts. The most derived Pachón cave population (Herman et al., 2018) is also starvation resistant but showed a nonsignificant change in symmetric NOA and DIR, perhaps because these cavefish have optimized their foraging under food-sparse conditions over the course of evolution (see below). In the Supplementary file 1—extended results and discussion, we also discuss age-dependent L–R plasticity in Tinaja cavefish (c.f., Figure 3B and C; Figure 5A and B) and the VAB levels of Los Sabinos and Tinaja cavefish.

### Advantages of laterality in the cave ecosystem

To understand the evolution of laterality and its plasticity in the context of cave ecology, we propose a model that explains the potential advantages of such lateralities and its plasticity in terms of calorie gain and consumption in food-sparse caves (Figure 6). This model assumes that (1) only one side of SNs accounts for VAB evocation and (2) starvation induces L–R symmetric NOA. Based on these assumptions, we hypothesized that sensory laterality and non-L–R-biased (i.e., symmetrical) approaches are beneficial in food-limited environments at the population level due to the tradeoff between calorie gain and calorie use in foraging.

**Figure 6.**
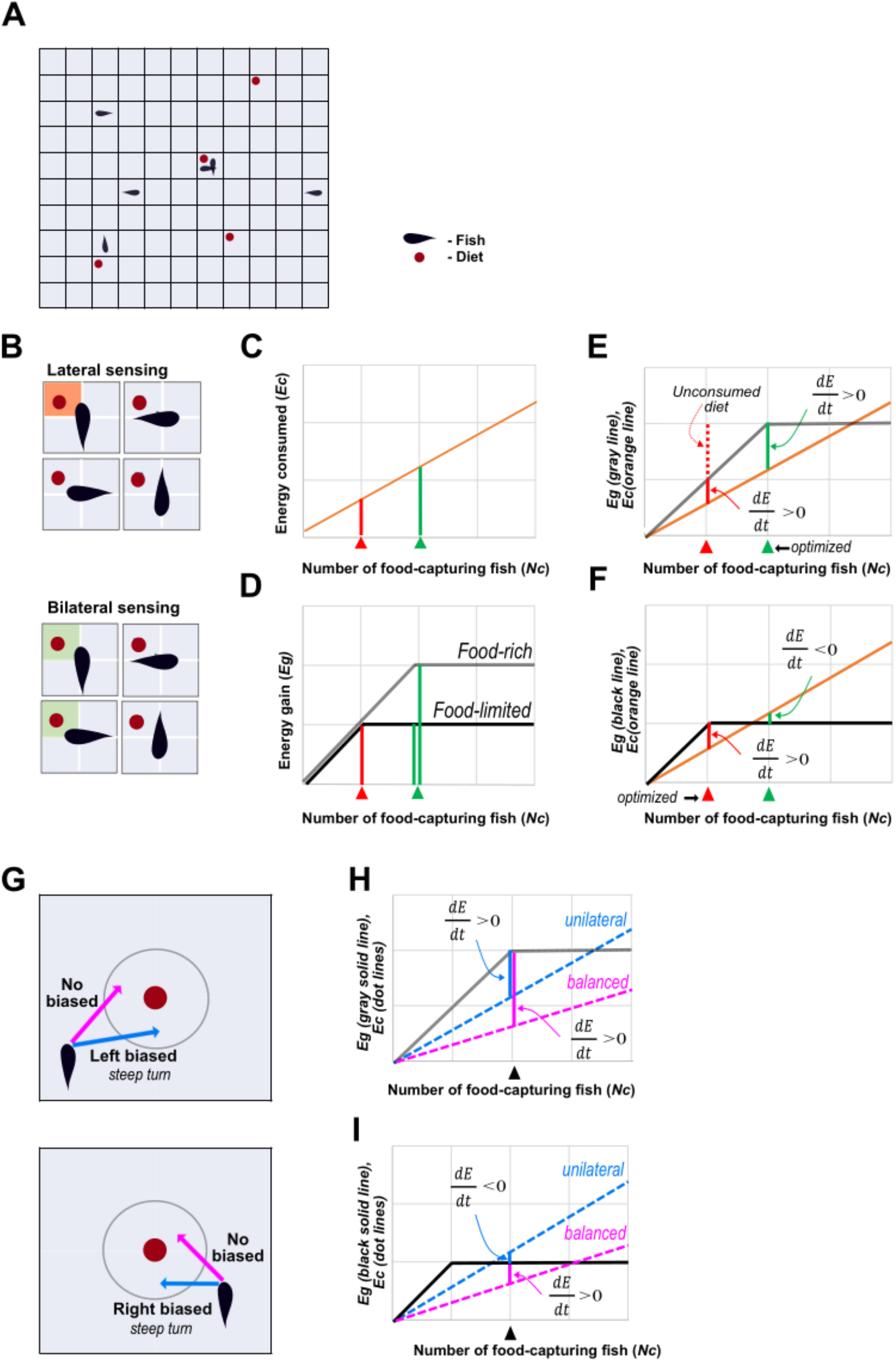
Relevance of sensory and behavior laterality. (**A**) General schematic of the model showing food availability and the number of fish. Each cell represents the sensing range of a fish. Red dots represent diet and black shapes represent fish. (**B**) In a given cell, the food detection probability *p_d_* is 1/4 for lateral sensing (top) and 1/2 for bilateral sensing (bottom); red and green shaded areas represent overlapping areas in the food and sensing fields, respectively. (**C**) Energy consumption (*E_c_*) as a function of the number of food-acquiring fish (*N_c_*). (**D**) Energy gain from food (*E_g_*) as a function of the number of food-acquiring fish (*N_c_*). In **C** and **D**, red arrows and lines indicate lateral sensing fish and green arrows and lines indicate bilateral sensing fish. Gray and black lines are the functions *E_g_* and depend on food-rich and - limited conditions, respectively. (**E, F**) Evaluation of 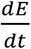. **E** shows *E_g_* and *E_c_* under food-rich conditions, whereas **F** shows *E_g_* and *E_c_* under food-limited conditions. 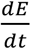 is based on the subtraction of *E_c_* from *E_g_* and is shown as solid vertical lines (green and red). In **E** (food-rich condition), the bilateral population is optimized because 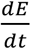 of the bilateral sensing population is larger than that of the unilateral sensing population. Furthermore, there is no unconsumed food in the bilateral sensing population, whereas unconsumed food is present in the unilateral sensing population (red dotted line). In **F** (food-limited condition), the unilateral population is optimized because 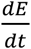 of the unilateral sensing population has a positive value, whereas 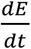 in the bilateral sensing population is negative. (**G**) Food-approaching behavior. The gray circle shows the 1.3-cm radius from the food (red circle). Blue and pink arrows represent steeper and less steep turns, respectively. Fish with balanced approaches take either left or right approaches, whereas fish with biased approaches (laterality) always take one-sided approaches that frequently require steeper turns. (**H, I**) Evaluation of 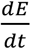 in food-rich (**H**) and food-limited (**I**) environments. The *E_c_* values for unilateral (steeper turn) fish and balanced fish are shown as blue and pink dotted lines, respectively. 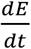 is based on the subtraction of *E_c_* (dotted lines) from *E_g_* (gray solid line in H or black in I); therefore, it is expressed as the blue and pink vertical solid lines. Under food-rich conditions, 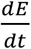 for both unilateral and balanced fish is positive; thus, both populations can gain energy. In contrast, under food-limited conditions, the L–R balanced population is optimal because it shows a positive value for 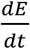. See Supplementary file 1 for further information.

The model, including food/energy gain, energy loss by thrasting, and the change in the total amount of calories gained in a population (= a sum of fish individuals), can be described as the following function:

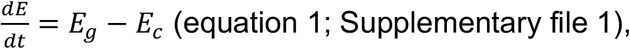

where *E* represents the total calories/energy of an average fish in a population gained per unit of time *t* (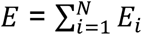, where *i* represents each individual fish and *E_t_* represents the total calories each fish gains per unit time), *E_g_* represents the average total calories the population gains from the diet per unit of time, and *E_c_* is the average total calories the population uses during foraging per unit of time.

Food capturing movements are assumed to use more calories, as the fish perform rapid and short turns toward their food, than are used in a typical swimming motion (personal observation). If the energy gain, *E_g_*, is higher than the energy consumption, *E_c_* (eq. 1), the current fish in the population can survive in the given environment. If the opposite is true (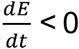; eq. 1), the number of fish in the population is reduced. We expect that unilateral sensing has an advantage over bilateral sensing in terms of the tradeoff between *E_g_* and *E_c_*.

Depending on two scenarios, i.e., (1) only a unilateral input or (2) the inputs of both sides are accounted for in VAB expression, the probability of sensing the food (*p_d_*) is defined as follows:

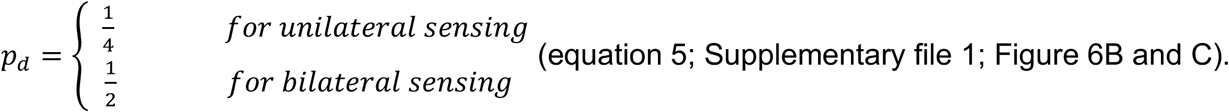

When a diet is available in a given cell area and two fish exist in the same cell area, two possibilities exist (Figure 6A): (1) in unilateral sensing, only one fish can sense and acquire the diet 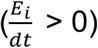, whereas the other fish does not evoke VAB, i.e., no calories are lost in the other fish 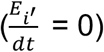 (Figure 6A and B); (2) in bilateral sensing, the two individuals compete for the diet unit (which only supplies one fish), resulting in one fish capturing the diet and gaining a positive *E* value 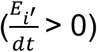 and the other losing calories due to their foraging activity 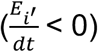, which results in many fish losing energy in the population (Figure 6C). The total 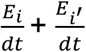 in unilateral sensing (1) is larger than that in bilateral sensing (2). This example shows that the number of fish reacting to and capturing food will differ in bilateral- or unilateral sensing situations, resulting in differences in *E* based on *E_c_* in equation 2 (Supplementary file 1): *E_c_* (calories consumed) depends on *p_d_* (eqs. 2, 4, and 5; Supplementary file 1), so the *E_c_* for bilateral sensing is twice as high as that for unilateral sensing (Figure 6C). As shown in Figure 6D and E, in a diet-rich environment, the bilateral sensing population has an advantage because its calorie gain is higher than that of the unilateral sensing population. Under food-sparse conditions, the advantage is reversed because the unilateral sensing population maintains 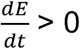, whereas the bilateral sensing population does not achieve 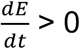 because many failed foraging attempts increase *E_c_* (Figure 6D and F). In summary, the unilateral sensing population can survive better than the bilateral sensing population by reducing the number of failed foraging attempts in food-limited conditions. This model suggests that the sensory laterality of Pachón cavefish could be beneficial in food-limited environments. The laterality in approaches (NOA) is defined as a preference for using the left or right-side of the head/body when approaching the vibration source. Our video observations indicated that fish exhibit multiple quick turns against the vibrating rod when they are in its vicinity (within a 1.3-cm radius of the rod). We assume that sharp turns cost more calories than typical swimming. We also assume that the fish that use unilateral approaches make steeper turns more often than the fish that use bilateral approaches (Figure 6G). The different levels of calorie consumption in fish using unilateral and bilateral approaches are implemented in the parameter ‘α’, the coefficient of *E_c_*, i.e., the average calorie consumption per time unit (eq. 2 in Supplementary file 1). Accordingly, the parameter *‘a’* value of the fish that use unilateral approaches is higher than that of the fish that use bilateral approaches (hatched lines in Figure 6H and I; Supplementary file 1). With this setting, under food-rich conditions, the 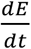 values of fish that use either unilateral or bilateral approaches are positive (Figure 6H); thus, both types of fish are expected to survive. In contrast, a population that uses bilateral approaches has a higher chance of survival 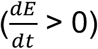 than a population that uses unilateral approaches in a food-limited environment (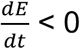; Figure 6I). Our starvation experiment showed that Tinaja cavefish shifted their approach from unilateral to L–R balanced (Figure 5A and 5C); therefore, these fish seem to adjust their L–R approach depending on food availability.

According to our model, Pachón cavefish is the most optimized population as it shows L–R symmetrical approaches (Figure 5A and 5C) as well as left-biased unilateral sensing (Figs. 1 and 2). The degree of laterality in Pachón and Tinaja cavefish populations fits well with their ecological conditions where Pachón cavefish experiences seasonal fluctuation of animated diets, crustaceans, which seems to be available more in the ~6-month dry season than in the 6-month wet season (Espinasa et al., 2017; Mitchell et al., 1977). In contrast, several Tinaja cave pools receive percolating water most of the time and have thick organic matter covering the bottom, distinct from Pachón cave, and the plastic laterality may be sufficient for adaptation to cave environment (Mitchell et al., 1977). There might be a possibility that Pachón cavefish reside in a more diet-limited condition than Tinaja, yielding further diversified laterality to adapt to cave environment.

In summary, laterality in sensory–behavior coupling and symmetrical approaches may have advantages in food-sparse environments and may have promoted adaptation processes in Pachón cave. Over the course of evolution, Tinaja cavefish evolved a plastic foraging strategy according to food availability, whereas Pachón cavefish evolved a robust SN sensory laterality with balanced approaches to vibrating objects. These results represent new insights that help us better understand the evolution of laterality, and they may lead to further insights into the diversification of laterality among animals.

## Materials and Methods

### *Astyanax mexicanus* husbandry and ethics statement

The lab-raised surface fish population used in this study is descended from an original collection from Balmorhea Springs State Park (Texas, USA), whereas the lab-raised cave populations used in this study are descendants of an original collection from Cueva de El Pachón in Tamaulipas collected by Richard Burowski (2009/2011), and collections from Sótano de la Tinaja and Cueva de Los Sabinos in San Luis Potosí, Mexico collected by Bill Jeffery in (2000 - 2001). The adult fish (1–3 years old) used in the experiments were raised in groups of 20–25 individuals in 0.74-L Ziploc containers (S.C. Johnson & Son Inc., Racine, WI, USA) and 1.4-L Zebrafish Aquatic Housing System tanks (Aquaneering Inc., San Diego, CA, USA). Fish were fed twice per day *ad libitum* with live *Artemia salina* larvae (Brine Shrimp Direct, Ogden, UT, USA).

All animal procedures were performed under protocol 17-2560 approved by the Institutional Animal Care and Use Committee at the University of Hawai‘i.

### Recording VAB

VAB assays were performed as described previously by Yoshizawa et al. (2010) and Worsham et al. (2019). Briefly, fish were acclimated for 4–5 days in a cylindrical chamber (10 cm in diameter; 5 cm in height; Pyrex 325 mL; Corning, NY, USA) filled with conditioned water (pH 6.8–7.0; conductivity of ~700 μS). Fish were fed once per day *ad libitum* with live *A. salina* larvae. After acclimation, the behavior of fish in response to vibrations was recorded for 3 min immediately after the introduction of the vibration rod into the bowl. The vibration stimulus was generated using a 7.5-mm diameter glass rod at 40 Hz with a GW Instek SFG-1003 DDS Function Generator (Good Will Instrument Co., Ltd., New Taipei City, Taiwan) connected to an audio speaker (ProSpeakers, Apple, Cupertino, CA, USA). Video footage of behavior was recorded under infrared illumination (880-nm wavelength; BL41192-880 blacklight; Advanced Illumination, Rochester, VT, USA) using a customized Microsoft MAIN-31891 LifeCam Studio installed with a Zoom lens (Zoom 7000, Navitar, Rochester, NY) and VirtualDub2 video capture software (licensed under the GNU General Public License: https://sourceforge.net/projects/vdfiltermod/).

The NOA was analyzed using ImageJ macro plugins (ImageJ 1.53a; NIH, Bethesda, MD, USA) and a homemade script published previously (Worsham et al., 2019; Yoshizawa et al., 2010). Using this method, we measured NOA and selected Pachón cavefish that exhibited NOA ≥ 6 for the ablation study (see “Ablation experiments” below).

### VAB video analysis using DLC

Videos were analyzed using the DLC toolbox version 2.0.6 (Mathis et al., 2018; Nath et al., 2019) in Python 3.6. DLC provides a robust suite of tools for deep learning-based markerless pose estimation. The approach is based on transfer learning, wherein a deep neural network is initially trained on an initial task with large amounts of available training data to achieve robust initial parameterization and then fine-tuned on a target task (ideally as closely related to the initial task as possible; in this case, pose estimation) using a smaller dataset. DLC uses a ResNet50 (He et al., 2016) backbone pretrained on the ImageNet (Deng et al., 2009) dataset to classify 1000 object categories in a dataset of over one million hand-labeled images. This network parameterization is then transferred and fine-tuned, during which the last layer of the network used originally for classification is replaced with transposed convolutional layers for pose estimation. These new layers predict individual probability densities over all pixels of a given input image for each pose marker, and such predictions are repeated independently for each frame in the video. DLC was also used to simplify data selection, data labeling, training, evaluation, and retraining processes. In this study, we specified a pose estimation task to track both the fish and rods in the VAB videos. For the fish, three head markers were tracked (right-side, left-side, and center) as well as one marker in the caudal fin; additionally, one marker was placed in the center of the vibrating rod. Because some videos contained two Petri dishes and some contained one, two separate sets of the same markers were allocated. The initial training set included frames selected randomly from 88 different VAB videos, and 807 hand-labeled training frames (images) were used to train our final model. The ResNet50 model was fine-tuned until the DLC early-stopping criterion was reached while using the default DLC training parameters as described by Mathis et al. (2018). All weights in the network were adjusted during training. The trained model was used to predict marker locations in all videos, outputting the x and y coordinates associated with the highest probability pixel for each marker. In frames with at least one DLC marker below a predefined confidence threshold, all marker locations were interpolated based on the markers’ positions in the preceding and following frames, which ensured that fish tracking was smooth throughout the duration of each video. Specifically, for a sequence of one or more subsequent frames where each frame had at least one marker below the confidence threshold, all markers’ locations were linearly interpolated between each of the respective markers’ locations in the frame immediately preceding and the frame immediately following that sequence. For videos including multiple recording arenas, markers for each Petri dish were treated independently. The confidence threshold was set empirically to 0.92 after manual inspection of the DLC outputs. The resulting marker locations were used to track several behaviors of interest in the videos: time spent within a 1.3-cm radius of the rod (DIR), time spent being outside of this radius (DOR), left- and right-sided approaches (NOA), and total swimming distance. The 1.3-cm radius from the vibrating rod was determined as the cutoff of VAB because it was the distance that best discriminated between the surface fish and cavefish approaches across the tested radii of 0.5– 2.5 cm (in 0.1-cm increments) from the vibrating rod. A left-sided approach was defined as entering the 1.3-cm radius threshold with the head-center marker while the head-left marker was closer than the head-right marker to the rod, and vice versa for a right-sided approach (Figure 3A). Accordingly, DIR, DOR, and NOA were separated as right- or left-facing based on the proximity of the head-right and head-left markers to the rod (Figure 3A).

### Repeatablity test

Repeatablity test was performed as described previously (Yoshizawa et al., 2010). Briefly, we first acclimated 22 surface, 9 Pachón cave and 11 Tinaja cave fish in cylindrical chambers for 4–5 days. Fish were fed once per day *ad libitum* with live *A. salina* larvae and recorded their movement as described in the Recording VAB section. After the recording, the cylindrical chambers were cleaned and fish were maintained in the same chambers for 6 days, and rerecorded in 7^th^ days of the second acclimation (6 days a part). This procedure was repeated for the 3^rd^ recording. The conditioned water was replaced every two days during this period. After the behavior analysis using DLC, interclass correlation repeatability (3 repeated measurements) was tested as referring Single_random_raters by the ‘psych’ R package.

### SN imaging

Neuromast vital staining was performed as described previously (Jørgenson and Jørgensen, 1989; Worsham et al., 2019; Yoshizawa et al., 2010). Fish were immersed in 25 μg/mL of 4-(4- (dimethylaminostyryl)-1-methylpyridiniumiodide (4-Di-1-ASP; MilliporeSigma, Burlington, MA) dissolved in conditioned water for 40–60 min, after which they were anesthetized in 66.7 μg/mL of ice-cold ethyl 3-aminobenzoate methanesulfonate salt (MS222; MilliporeSigma) in conditioned water (pH 6.8–7.0; conductivity of ~700 μS). Fish were then visualized using a fluorescence microscope [BX61WI Olympus microscope with a 2.5× MPlanFL N lens, a Rhodamine filter set, and an ORCA-Flash 4.0 digital camera (Olympus Corp.)]. SNs were counted from digital images of 4-Di-1-ASP-stained fish using the “Analyze Particles” function of ImageJ 1.53a. SNs were counted both in the left and right sides of the third infraorbital/suborbital bone.

### Ablation experiments

Ablation experiments were performed using 1-year old surface fish (n = 47) and VAB-positive cavefish (Pachón population, n = 67). The VAB level was measured before ablation, and VAB-positive cavefish (NOA ≥ 6 according to ImageJ-based analysis) were selected strategically to determine the association between IO3 SNs and VAB level.

SN ablations were performed using Vetbond (3M, St. Paul, MN, USA), a nontoxic tissue adhesive, on the left (surface fish, n = 14; Pachón cavefish, n = 30), right (surface fish, n = 13; Pachón cavefish, n = 25), and both left and right (surface fish, n = 10; Pachón cavefish, n = 12) sides (Figure 2A and B). Fish were anesthetized in 66.7 μg/mL of ice-cold ethyl 3-aminobenzoate methanesulfonate salt (MS222, MilliporeSigma) in conditioned water (pH 6.8– 7.0; conductivity of ~700 μS). Excess water was removed carefully using Kimtech wipes, and the adhesive (0.5 μL) was applied to the skin surface. After being exposed to the air for 10 s, treated fish were placed in bowls (10 cm in diameter; 5 cm in height; Pyrex 325 mL, Corning) filled with conditioned water at room temperature. Within a day of active swimming, the adhesive usually peeled off, and VAB was assayed. 4-Di-1-ASP staining was performed to count the number of SNs in the IO3 region before and after ablation.

### Fasting

Pachón cavefish (n = 20), Tinaja cavefish (n = 20) and surface fish (n = 16) individuals (~1 year old) were placed in cylindrical chambers (10 cm in diameter; 5 cm in height; Pyrex 325 mL, Corning) and acclimated for 4–5 days in conditioned water (pH 6.8–7.0; conductivity of ~600 μS) in which they were fed once a day (weight or approximate number of *Artemia*) with live *A. salina* larvae. After acclimation, VAB was recorded and analyzed. Following this first assessment, fish were starved for 6 days and VAB was recorded and analyzed once again.

### Statistical analysis

All statistical analyses were performed in RStudio 4.0.3 (RStudio, Boston, MA, USA). The R packages used included lme4, lmerTest, car, coin, yarrr, AICcmodavg, and ggpubr. Linear or generalized linear models were selected using Akaike’s information criterion function to identify the best fit models for analysis of the NOA and DIR in all experiments. Analysis of variance (ANOVA) was performed using generalized linear model fitting functions (glm or glmer in the lme4 package). Post-hoc tests were performed using the Wilcoxon signed-rank (paired) test followed by Holm’s multiple-test correction. Nonparametric Kendall’s t correlation was used to evaluate associations between neuromasts and behavior outputs. In the field experiments, the total number of turns within the circle were pooled for each cave pool, and the left-side turns vs. the right-side turns were tested by fitting a generalized linear mixed-effects model using a binomial distribution.

## Acknowledgement

We thank L Espinasa for constructive comments regarding insights on variation of laterarities in *Astyanax* cavefish. We also thank to H Yoshizawa for supporting the repeatability experiments and fish care assistance. We are grateful to H Hernandez, N Doeden, C Balaan, V Crystal, J Choi, L Lu, J Nguyen, K Lactaoen, M Worsham, J Kato, M Ito, R Balmilero-Unciano, E Doy, A Martinez, and D Mones for fish care assistance. We gratefully acknowledge support from the National Institute of Health (P20GM125508) to MY, Hawaii Community Foundation (18CON-90818) to MY.

## Author contribution

VFLF: designed and performed experiments, wrote original draft, edited the manuscript, performed all experiments, and analyzed results.

YG: designed the detection and analysis algorithm and helped with actual analysis related to DeepLabCut-based aanalysis, consulted the experimental procedure, and edited the manuscript.

MI: performed experiments, edited the manuscript, analyzed the results, and developed a mathematical model.

MY: designed the experiment, edited the manuscript,co-developed DeepLabCut-based analysis method and algorithm, and analyzed the results.

---

**Figure 2—figure supplement 1.**
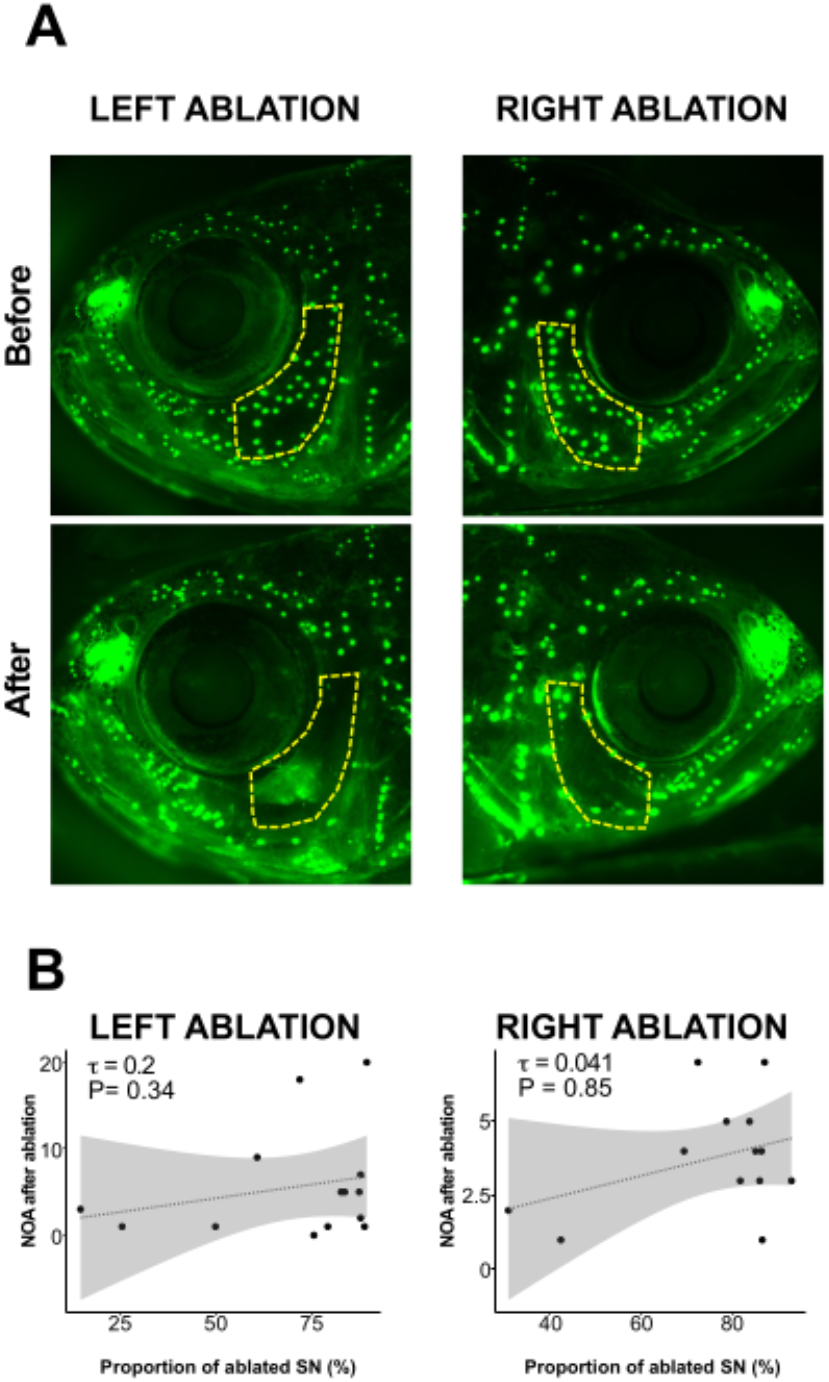
Ablation of superficial neuromasts (SNs) did not induce detectable effect on the level of VAB in surface fish. (**A**) Left and right sides of SNs in Pachón cavefish stained with 4-Di-1-ASP (green dots) before and after ablation of SNs in the infraorbital region (IO3 shown by a yellow line) of the same fish. (**B**) Proportion of ablated SNs on the left and right sides plotted against the total NOA in both cases. Regression lines with 95% confidence intervals (shaded gray or orange) are shown in each panel. See Supplementary Table 3 for additional details.

**Figure 2—figure supplement 2.**
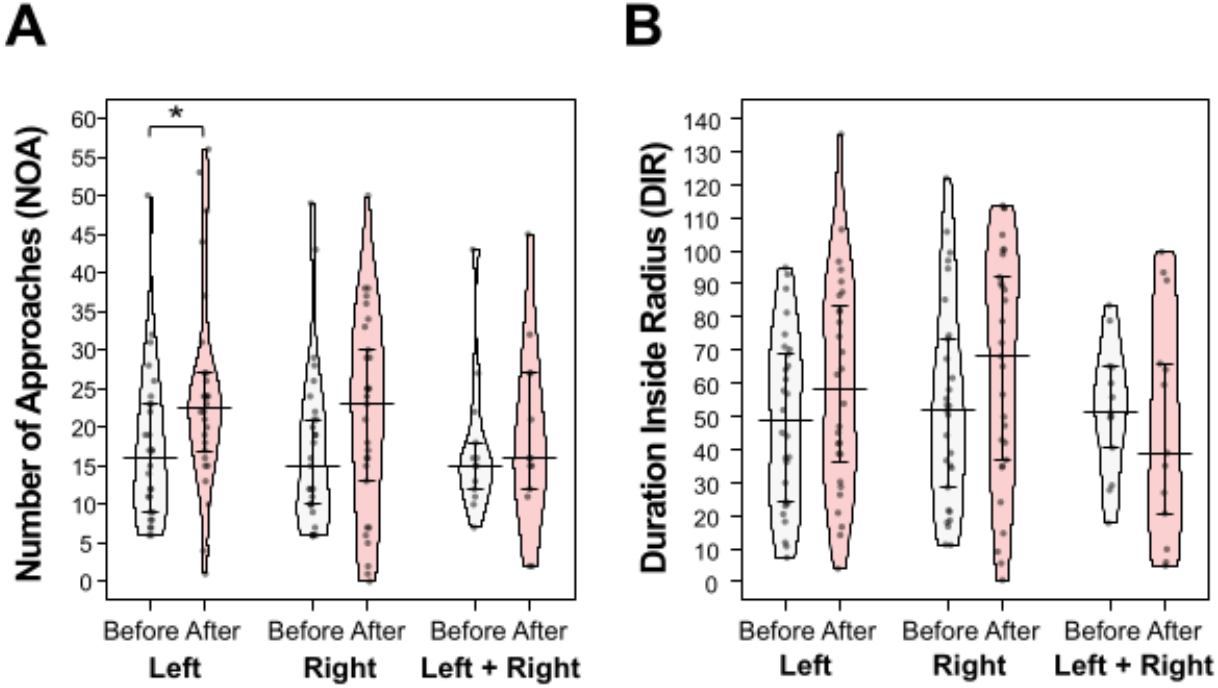
Overall results of the SN ablation experiments in Pachón cavefish. (**A**) Number of approaches (NOA) in Pachón cavefish. Ablation of the left SN resulted in an increase in NOA (V = 74.5, P = 0.032); however, ablation of the right SN changed NOA at an underdetection level (V = 121, P = 0.063). (**B**) Duratin in the radius (DIR) in Pachón cavefish. Left SN or right SN ablation did not show detectable changes in DIR. Pirate plot bars represent means ± standard errors of the means. Data were analyzed using a generalized linear model followed by a post-hoc test (Wilcoxon test, adjusted by Holm’s correction): *, P < 0.05; **, P < 0.01; and ***, P < 0.001. Statistical scores are available in Supplementary Table 4.

**Figure 3—figure supplement 1.**
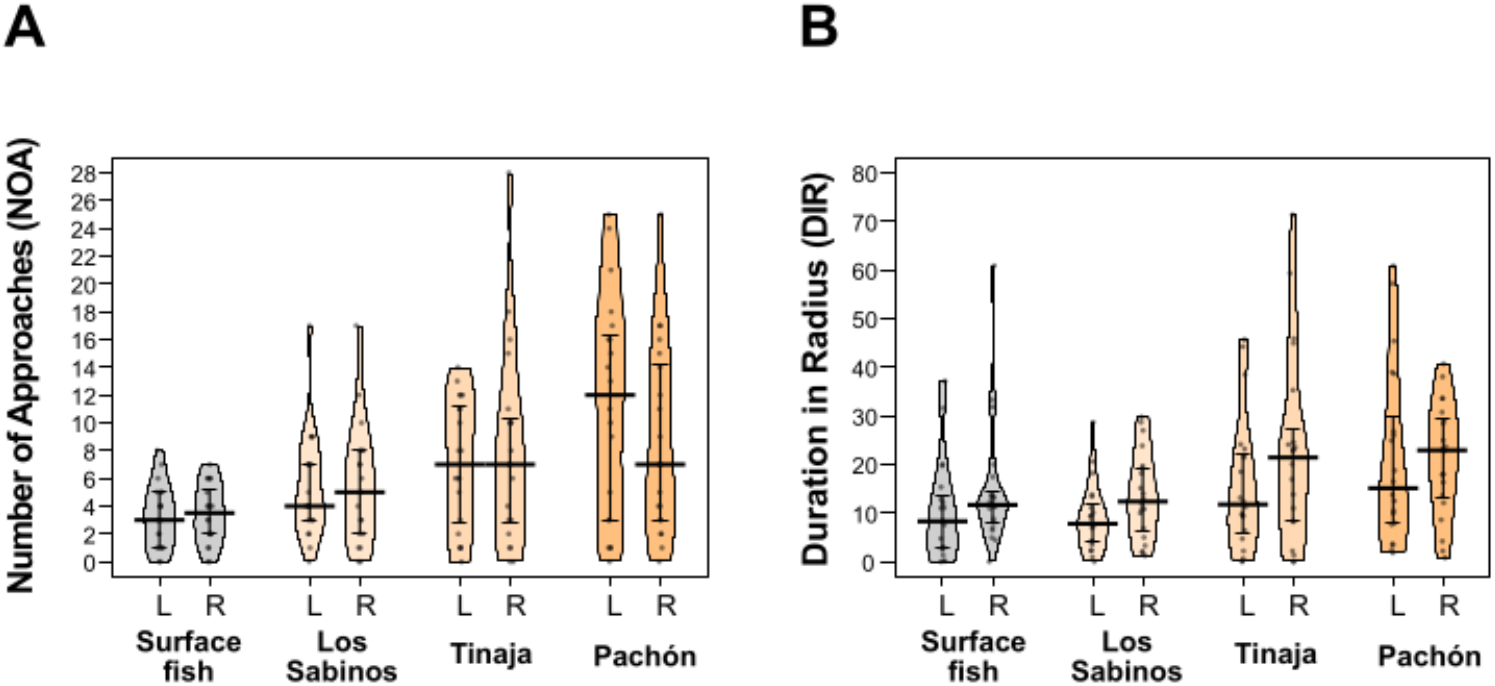
*Astyanax mexicanus* populations did not exhibit left- or right-biased approaches (NOA) or adhesion (DIR) at the population level. (**A, B**) Pirate plots showing no detectable differences between the number of left- and right-side approaches (NOA) (**A**) or the duration in the radius (DIR) (**B**) among the four tested populations (2–3-year-old individuals). Data represent means ± standard errors of the means (each population, n = 20). Further details are presented in Supplementary Table 1.

**Figure 5—figure supplement 1.**
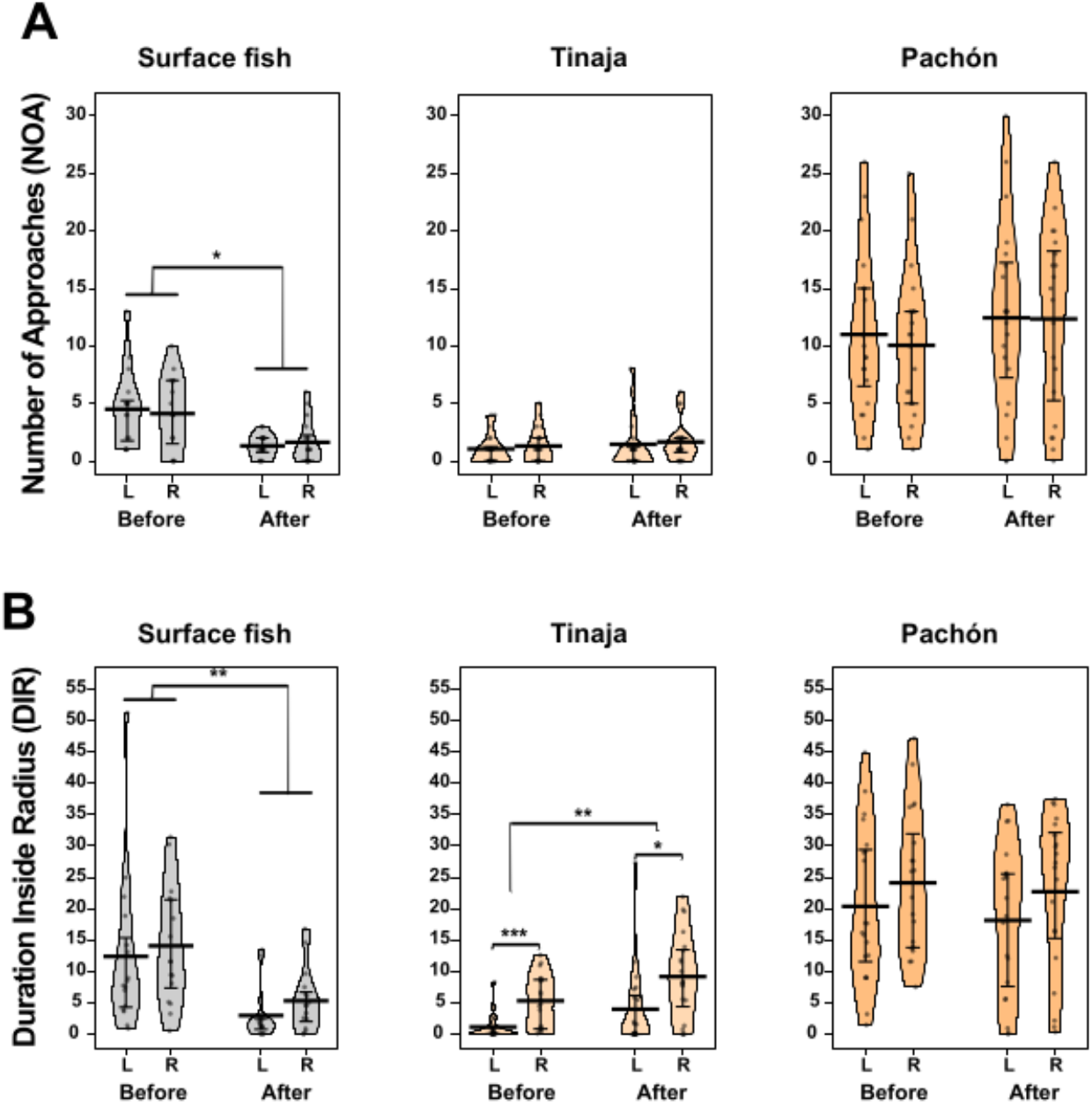
In response to starvation, lateralities in NOA and DIR are plastic in Tinaja cavefish but not in Pachón cavefish. (**A, B**) Pirate plots showing the number of left- and right-side approaches (NOA) (**A**) and left- and right-side-associated durations within a 1.3-cm radius (DIR) (**B**) before and after a 6-day fasting period. In **A**, only surface fish showed a significantly reduced NOA after fasting among the three tested populations, and no L–R bias (laterality) was detectable. In **B**, surface fish showed significantly reduced DIR after fasting, whereas Tinaja cavefish showed increased DIR after fasting. Tinaja cavefish also showed right bias in DIR before and after fasting, but they showed more significant right bias than left bias in DIR before fasting. Pachón cavefish did not exhibit any changes in DIR or L–R bias in DIR. All statistical scores are available in Supplementary Table 5. *, P<0.05; **, P<0.01.

