## Supplementary file 1 and Tables for "Possible role of left–right asymmetry in the sensory system and behavior during adaptation to food-sparse cave environments"

**Supplementary file 1**
**Extended results and discussion**
*VAB of Tinaja cavefish was higher than that of Los Sabinos cavefish, contradicting results reported previously.*
In Figure 1C, Tinaja cavefish are shown to have a similar VAB level to that of Pachón cavefish, both in terms of NOA (Pachón vs. Tinaja: P = 0.620) and DIR (Pachón vs. Tinaja: P = 0.232) (also see Supplementary Table 1). Contrastingly, the VAB level of Los Sabinos cavefish (both NOA and DIR) was indistinguishable from that of surface fish (surface fish vs. Los Sabinos: NOA, P = 0.054; DIR, P = 0.795; Figure 1C and Supplementary Table 1). In a previous study, Los Sabinos cavefish showed a higher VAB level than that of Tinaja cavefish (Yoshizawa et al., 2015). Thus, VAB may be plastic in Tinaja and Los Sabinos cavefish. Indeed, we detected a plastic change in VAB in Tinaja cavefish according to food availability and age (see the main text). Interestingly, Tinaja and Los Sabinos caves are close geographically and thought to be connected (Mitchell et al., 1977).

*Increase in NOA after SN ablation in Pachón cavefish*
In Pachón cavefish, ablation of the left SNs (but not right SNs or left and right SNS) induced an increase in NOA (Figure 2—figure supplement 1). The increase in NOA after SN ablation contradicted the findings of previous studies in which SN ablation (both left and right SNs) reduced NOA (Yoshizawa et al., 2010, 2012). These discrepant results may be due to the following: (1) the selection of Pachón cavefish individuals with high VAB levels (NOA ≥ 6; see Materials and Methods) for ablation in the present study, potentially including individuals with high appetite/internal motivation by default; (2) the use of an SN ablation area in the IO3 area that was not sufficiently large at this stage in the fish (i.e., a larger SN ablation area in the trunk reduces NOA; Yoshizawa et al., 2010); (3) an inflammatory reaction after injury that enhanced sensitivity (Neumann et al., 1996; Wang et al., 2004); and (4) individual Pachón cavefish with fewer left SNs showing plasticity in NOA (increased NOA) in response to fasting (Figure 4C). In the current study, we did not test any of these possibilities; however, considering that one or more of these explanations is accurate, the tested fish might express a higher response to the vibration stimulus even with the loss of SNs. These possibilities should be investigated further in subsequent research.

*Laterality in Tinaja cavefish according to age*
In the lab, 1-year-old Tinaja cavefish showed a right-side bias in DIR before and after starvation (Figure 5—figure supplement 1A). Thus, 1-year-old Tinaja cavefish with right-side bias showed plasticity in DIR in response to food availability. In addition, the degrees of L–R bias in NOA and DIR were also plastic and became symmetrical after starvation (Figure 5A, B; Figure5—figure supplement 1).
Interestingly, the right-side bias found in 1-year-old Tinaja cavefish was not detected in 2–3-year-old Tinaja cavefish. Notably, the overall VAB level (NOA and DIR) observed in 1-year-old Tinaja cavefish was lower than that in 2–3-year-old Tinaja cavefish (Figure 3—figure supplement 1; and Figure 5—figure supplement 1). Previous studies have shown that VAB level depends on age in Pachón cavefish (Yoshizawa et al., 2010, 2012) and varies among pools within the same cave (Pachón, Tinaja, and Molino caves) (Espinasa et al., 2021). Considering these findings, L–R bias in NOA and DIR and the overall levels of NOA and DIR could depend on environmental conditions and developmental stage. However, given standardized conditions, these L–R biases and VAB levels are consistent in the same individuals (Yoshizawa et al., 2010; also see the main text).
In summary, the L–R bias in VAB and overall VAB levels can vary even within the same cave population. Among the four populations tested here, Tinaja cavefish were the most plastic in terms of VAB level and L–R bias.

*Relevance of sensory and behavior laterality in terms of calorie gain and consumption*

1. Sensory laterality

We hypothesized that sensory laterality is beneficial in a food-limited environment in terms of the tradeoff between calorie gain and calorie consumption at the population level.

The change in the total amount of calories held in a population can be described as follows:

$$\frac{dE}{dt} = E_g - E_c, \tag{1}$$

where $E$ is total calories (energy), $t$ is time, $E_g$ is the average total calorie gain due to food uptake per unit time, and $E_c$ is the average total calorie consumption per population per unit time due to food capturing (foraging). Compared with the typical swimming motion, food capturing motions are assumed to consume more calories because they involve quick turns toward the food item (personal observation of how the studied fish acquire food aggressively). When $\frac{dE}{dt} > 0$, the population can survive in a given environment; when $\frac{dE}{dt} < 0$, the number of individuals in a population will decline.

We expect that sensory laterality is a key property affecting the tradeoff between $E_g$ and $E_c$.

To deliver $E_g$ and $E_c$, we assume the following conditions:

| Assumption | Description |
|---|---|
| **1** | A small cell area is defined as the range area in which the fish can sense food (Figure 6A). The whole area is defined as *A,* i.e., the number of cell areas in the whole area. |
| **2** | The amount of food supplied in a unit of time in a field (denoted as $F$) is constant, and in the next time step the food is gone, i.e., the number of food items in a field is always same. |
| **3** | Food items are randomly dispersed in a field; however, for simplicity, food items are always in the top left corner in a cell area. |
| **4** | Fish exist in random positions and directions in an entire field. For simplicity, fish are positioned at the center of the cell area. |
| **5** | The number of food items captured by a fish is set as 1 (constant) when the food is in the same cell area. Even if there are two food items in the same cell area, the fish can only acquire one food item at a time. |

Thus, total calorie consumption per unit time, $E_c$, is given as follows:

$$E_c = a \times A \times p_f \times N_c, \tag{2}$$

where $a$ is the coefficient of the average calorie consumption per fish to reach the food in a cell, $A$ is the number of cells in the whole field, $p_f$ is the probability that food exists in a cell area, and $N_c$ is the number of fish engaging in food capturing in a cell area. Additionally, $p_f$ is calculated as follows:

$$p_f = \begin{cases} F/A \; if \; F < A \\ 1 \; if \; F \geq A \end{cases}. \tag{3}$$

$N_c$ is calculated as follows:

$$N_c = p_d \times \frac{N}{A}, \tag{4}$$

where $p_d$ is the probability of sensing the food when the food exists in a cell area. Depending on whether VAB is regulated by the unilateral or bilateral sensory input, the value of $p_d$ is defined as shown in Figure 6B.

$$p_d = \begin{cases} 1/4 \; for \; lateral \; sensing \\ 1/2 \; for \; bilateral \; sensing \end{cases}. \tag{5}$$

Note that the $N_c$ for unilateral sensing is half that for bilateral sensing. When only one food item in the cell area is available but two fish exist in the same cell area, the following apply: (i) in case of bilateral sensing, the two fish compete for one food item, resulting in one fish losing calories by failing to capture the food; (ii) for unilateral sensing, only one fish senses the food and acquires it, whereas the other fish does not evoke VAB and loses no calories.

As shown in eq. (2), $E_c$ is a linear function of $N_c$ (Figure 6C). Because the $N_c$ for bilateral sensing is twice that for unilateral sensing, the $E_c$ for bilateral sensing is also twice as high as that for unilateral sensing. Again, when multiple fish compete for limited food, we expect that the unilateral sensing population will save relatively more calories; therefore, such sensing can be considered adaptive to the environment because of reduced calorie consumption ($E_c$).

Total calorie gain by food uptake per unit time, $E_g$, is defined as follows:

$$E_g = b \times A \times F_C, \tag{6}$$

where $b$ is the coefficient of the efficacy of calorie gain from a food item and $F_c$ is the number of food items consumed in a cell area. $F_c$ is a function of $N_c$. If there are sufficient food items, $F_c$ increases linearly depending on how many fish are in a cell area. When food is limited, $F_c$ is saturated to the upper limit of food items in a cell. Thus, $F_c$ can be expressed using the following equations based on the assumption (Assumption 5) stated above (the number of food items a fish can consume per unit time $dt$ is 1):

$$F_c = \begin{cases} F/A \times N_c \;\; if \; F/A > N_c \\ F/A \; if \; F/A \le N_c \end{cases}. \tag{7}$$

In eqs. (6) and (7), $E_g$ is expressed as a function of $N_c$ in Figure 6D. As shown in Figure 6D, $E_g$ can differ depending on the food supply in an area. If the number of food items is large, $E_g$ has a linear relationship with $N_c$ even if the value of $N_c$ is large, but if the number of food items is small, $E_g$ is saturated to the upper limit.

Finally, $\frac{dE}{dt}$ in eq. (1) can be rewritten with eqs. (2) and (6) as a function of $N_c$ as follows:

$$\frac{dE(N_c)}{dt} = b \times A \times F_c(N_c) - a \times A \times p_f \times N_c, \tag{8}$$

where the parameters $a, b, A, \text{and } F$ have the following condition:

$$b \times F > a \times A. \tag{9}$$

Thus, eq. (8) must output a positive value at a certain domain, otherwise $\frac{dE(N_c)}{dt}$ is negative for every $N_c$ and the population size decreases.

Equation (8) can be examined by overlaying the graphs shown in Figure 6C and D, as shown in Figure 6E and F. $\frac{dE(N_c)}{dt}$ differs depending on whether the fish are in a food-rich or -limited environment. Under food-rich conditions, the bilateral sensing population is optimized because $\frac{dE(N_c)}{dt}$ is larger than that in the lateral sensing population (Figure 6E). Under food-limited conditions, however, unilateral sensing is optimized because $\frac{dE(N_c)}{dt}$ is positive, whereas that for bilateral sensing is negative (Figure 6F). In the Pachón cave, food is known to be limited (Espinasa et al., 2017); therefore, lateral sensing is expected to be optimal in the Pachón environment.

2. Laterality in approaches (behavior laterality)
Behavior laterality is defined as foraging approaches made dominantly from one side (left or right) after VAB is triggered in the fish. From video observations, fish were found to make sharp turns toward the vibrating rod when approaching in the vicinity of the rod (within 1.3 cm). We predict that sharp turning consumes more calories than typical swimming. We also assume that fish that show unilateral approaches turn with steeper angles compared with fish approaching the food from both sides. This assumption is based on the notion that the position of the fish relative to the food is stochastic (Figure 6G).

Here, the larger energy loss in the biased approaches is expressed as $E_c$ in eq. (1). Specifically, the difference between the unilateral or balanced approaches is expressed by the parameter $a$, the coefficient of the average calorie consumption per fish. Accordingly, $a$ has a higher value in fish that make unilateral approaches relative to those that make balanced approaches. These scenarios are shown in Figure 6H and 6I.

The value of $\frac{dE(N_c)}{dt}$ differs depending on whether conditions are food-rich or -limited. As shown in Figure 6H, $\frac{dE(N_c)}{dt}$ is positive for fish that make unilateral and balanced approaches under food-rich conditions; thus, both populations will gain energy. In contrast, under food-limited conditions, making balanced approaches is optimal because $\frac{dE(N_c)}{dt}$ has a positive value (Figure 6I). Our fasting experiment showed that Tinaja cavefish shifted their approach from unilateral to balanced; therefore, these cavefish are able to adjust their foraging behavior depending on food availability. The sensory-foraging behavior of Pachón cavefish is further optimized to food-limited conditions in terms of both sensing food (sensory unilaterality) and foraging behavior (balanced approach) because both unilateral sensing and balanced approaches can reduce wasting energy.

#### Supplementary Table 1. Statistical scores for Figs. 1 and 3: figure supplement 1.

| | | Generalized linear model Statistical score | P-values | *Wilcoxon* *(pairwise comparison)* | P-values *(Holm's correction)* |
|---|---|---|---|---|---|
| **Swimming Distance** | Population (SF, LS, TI, PA) | F(3, 76) = 14.17 | **2.00E-07*** ** | | |
| **Number of Approaches (NOA)** | Population (SF, LS, TI, PA) | F(3, 152)= 11.327 | **9.52E-07*** ** | SF vs LS | 0.054 |
| | | | | SF vs TI | **0.004** ** |
| | LR (Left, Right) | F(1, 152)= 0.149 | 0.700 | SF vs PA | **7.7E-04*** ** |
| | | | | LS vs TI | 0.232 |
| | Pop × LR | F(3, 152)= 0.754 | 0.522 | LS vs PA | 0.053 |
| | | | | TI vs PA | 0.232 |
| **Duration Spent Inside the Radius (DIR)** | Population (SF, LS, TI, PA) | F(3, 152)= 5.098 | **2.17E-03** ** | SF vs LS | 0.795 |
| | | | | SF vs TI | 0.282 |
| | LR (Left, Right) | F(1, 152)= 3.378 | 0.068 | SF vs PA | **0.026*** |
| | | | | LS vs TI | 0.282 |
| | Pop × LR | F(3, 152)= 0.498 | 0.684 | LS vs PA | **0.016*** |
| | | | | TI vs PA | 0.620 |
| **Duration Spent Outside the Radius (DOR)** | Population (SF, LS, TI, PA) | F(3, 152)= 0.861 | 0.463 | | |
| | LR (Left, Right) | F(1, 152)= 0.560 | 0.455 | | |
| | Pop x LR | F(3, 152)= 0.016 | 0.997 | | |

Surface fish (SF, n = 20) and Los Sabinos (LS, n = 20), Tinaja (TI, n = 20), and Pachón (PA, n = 20) cavefish.
Linear or generalized linear models were selected using Akaike's information criterion function to determine the best fit model for analyzing swimming distance, NOA, DIR, and DOR.
***: Significant at alpha = 0.001; **: significant at alpha = 0.01; *: significant at alpha = 0.05.

#### Supplementary Table 2. Correlation analysis for Figure 1.

| | SURFACE FISH | | LOS SABINOS | | TINAJA | | PACHÓN | |
|---|---|---|---|---|---|---|---|---|
| | **Kendall Tau** | **P-value** | **Kendall Tau** | **P-value** | **Kendall Tau** | **P-value** | **Kendall Tau** | **P-value** |
| **Correlation against total NOA** | | | | | | | | |
| Left SN number | −0.145 | 0.420 | −0.147 | 0.377 | −0.145 | 0.379 | **0.329** | **0.047*** |
| Right SN number | −0.158 | 0.378 | −0.264 | 0.116 | −0.048 | 0.770 | 0.258 | 0.118 |
| Total SN number | −0.130 | 0.466 | −0.201 | 0.227 | −0.145 | 0.379 | 0.269 | 0.103 |
| **Correlation against total DIR** | | | | | | | | |
| Left SN number | −0.238 | 0.172 | −0.027 | 0.871 | −0.095 | 0.559 | 0.292 | 0.074 |
| Right SN number | −0.223 | 0.198 | −0.053 | 0.745 | 0.106 | 0.516 | 0.212 | 0.194 |
| Total SN number | −0.223 | 0.198 | −0.058 | 0.721 | −0.053 | 0.745 | 0.265 | 0.104 |
| **Correlation against left NOA** | | | | | | | | |
| Left SN number | −0.272 | 0.133 | −0.185 | 0.277 | 0.125 | 0.452 | 0.312 | 0.059 |
| Right SN number | **−0.388** | **0.031*** | −0.175 | 0.306 | 0.267 | 0.109 | 0.263 | 0.110 |
| **Correlation against right NOA** | | | | | | | | |
| Left SN number | 0.070 | 0.699 | −0.141 | 0.396 | −0.243 | 0.142 | 0.269 | 0.103 |
| Right SN number | 0.098 | 0.589 | −0.225 | 0.180 | −0.102 | 0.536 | 0.241 | 0.143 |
| **Correlation against left DIR** | | | | | | | | |
| Left SN number | −0.277 | 0.111 | 0.000 | 1.000 | −0.011 | 0.948 | **0.324** | **0.047*** |
| Right SN number | **−0.341** | **0.049*** | 0.027 | 0.871 | 0.254 | 0.119 | 0.201 | 0.217 |
| **Correlation against right DIR** | | | | | | | | |
| Left SN number | −0.185 | 0.288 | −0.090 | 0.581 | −0.133 | 0.417 | 0.245 | 0.135 |
| Right SN number | −0.144 | 0.404 | −0.086 | 0.602 | −0.037 | 0.820 | 0.164 | 0.314 |

ANOVA was performed using the generalized linear model fitting function. Population differences for NOA and DIR were determined via pairwise comparisons using a Wilcoxon rank sum test with continuity correction. Holm's correction method was used for P-value adjustment.
***: Significant at alpha = 0.001; **: significant at alpha = 0.01; *: significant at alpha = 0.05.

#### Supplementary Table 3. Correlation analysis for Figure 2.

| | Surface fish | | Pachón cavefish (number of approaches > 6) | |
|---|---|---|---|---|
| | **Kendall Tau R** | **P-value** | **Kendall Tau R** | **P-value** |
| **Correlation against Total NOA after ablation** | | | | |
| Left SN ablation (%) | 0.198 | 0.342 | **−0.342** | **0.009**** |
| Right SN ablation (%) | 0.041 | 0.852 | −0.064 | 0.635 |
| Left + Right SN ablation (%) | 0.159 | 0.528 | −0.237 | 0.269 |
| **Correlation against Total DIR after ablation** | | | | |
| Left SN ablation (%) | 0.155 | 0.443 | −0.094 | 0.479 |
| Right SN ablation (%) | 0.179 | 0.435 | 0.003 | 0.984 |
| Left + Right SN ablation (%) | 0.289 | 0.291 | −0.026 | 0.952 |
| **Correlation against Left NOA after ablation** | | | | |
| Left SN ablation (%) | 0.189 | 0.383 | −0.376 | **0.004**** |
| Right SN ablation (%) | 0.042 | 0.850 | −0.032 | 0.812 |
| **Correlation against Right NOA after ablation** | | | | |
| Left SN ablation (%) | 0.070 | 0.753 | −0.057 | 0.677 |
| Right SN ablation (%) | 0.128 | 0.539 | −0.189 | 0.148 |
| **Correlation against Left DIR after ablation** | | | | |
| Left SN ablation (%) | 0.287 | 0.154 | −0.147 | 0.254 |
| Right SN ablation (%) | 0.116 | 0.582 | −0.098 | 0.465 |
| **Correlation against Right DIR after ablation** | | | | |
| Left SN ablation (%) | −0.013 | 0.951 | 0.056 | 0.678 |
| Right SN ablation (%) | 0.122 | 0.546 | −0.085 | 0.524 |

***: Significant at alpha = 0.001; **: significant at alpha = 0.01; *: significant at alpha = 0.05.

Left SN: superficial neuromasts in the left-side; Right SN: superficial neuromasts in the right-side; Left + Right SN: sum of the number of superficial neuromasts present in both left and right sides; NOA: number of approaches; DIR: duration inside a 1.3-cm radius from the rod; Left NOA: number of approaches to the rod with the left-side of the head; Right NOA: number of approaches to the rod with the right-side of the head; Left DIR: duration inside a 1.3-cm radius from the rod with left-side head facing the rod; Right DIR: duration inside a 1.3-cm radius from the rod with right-side head facing the rod; Total NOA: sum of Left NOA and Right NOA; Total DIR: Sum of Left NOA and Right DIR.

Only Kendall Tau R values that satisfy P < 0.05 with 95% confidence intervals that do not cross 0 are shown in bold.

#### Supplementary Table 4. Statistical scores for Figure 2—figure supplement 2.

| | *ANOVA* | | | *Wilcoxon signed-rank (paired)* | | | *Holm's correction* | |
|---|---|---|---|---|---|---|---|---|
| | | **statistics** | **p-value** | | **statistics** | **p-value** | **alpha = 0.05** | **K = 3** |
| **Swimming Distance** | Ablation | $\chi^2(2)$= 0.7 | 0.965 | | | | | |
| ***(glm)*** | Prepost | $\chi^2(1)$= 1.3 | 0.245 | | | | | |
| | Ablation:Prepost | $\chi^2(2)$= 4.8 | 0.091 | | | | | |
| **Number Of Approaches** | Ablation | $\chi^2(2)$= 1.7 | 0.431 | PA Left (Pre vs. Post) | V = 74.5 | 0.011 | **0.032*** | |
| **(NOA)** | Prepost | $\chi^2(1)$= 8.8 | **0.003**** | PA Right (Pre vs. Post) | V = 121 | 0.063 | | |
| **(glm)** | LR | $\chi^2(1)$= 0.1 | 0.713 | PA Left+Right (Pre vs. Post) | V = 39 | 1 | | |
| | Ablation:Prepost | $\chi^2(2)$= 1.1 | 0.586 | | | | | |
| | Ablation:LR | $\chi^2(2)$= 0.1 | 0.973 | | | | | |
| | Prepost:LR | $\chi^2(1)$= 0.5 | 0.479 | | | | | |
| | Ablation:Prepost:LR | $\chi^2(2)$= 0.9 | 0.643 | | | | | |
| **Duration Spent Inside** | Ablation | $\chi^2(2)$= 2.6 | 0.276 | PA Left (Pre vs. Post) | V = 139 | 0.150 | | |
| **The Radius (DIR)** | Prepost | $\chi^2(1)$= 4.3 | **0.039*** | PA Right (Pre vs. Post) | V = 138 | 0.088 | | |
| **(lmer)** | LR | $\chi^2(1)$= 0.0 | 0.964 | PA Left+Right (Pre vs. Post) | V = 50 | 0.787 | | |
| | Ablation:Prepost | $\chi^2(2)$= 2.4 | 0.306 | | | | | |
| | Ablation:LR | $\chi^2(2)$= 0.3 | 0.858 | | | | | |
| | Prepost:LR | $\chi^2(1)$= 0.4 | 0.506 | | | | | |
| | Ablation:Prepost:LR | $\chi^2(2)$= 0.1 | 0.950 | | | | | |
| **Duration Spent Outside** | Ablation | $\chi^2(2)$= 0.4 | 0.800 | | | | | |
| **The Radius (DOR)** | Prepost | $\chi^2(1)$= 1.6 | 0.202 | | | | | |
| **(lmer)** | LR | $\chi^2(1)$= 3.6 | 0.058 | | | | | |
| | Ablation:Prepost | $\chi^2(2)$= 2.7 | 0.256 | | | | | |
| | Ablation:LR | $\chi^2(2)$= 4.1 | 0.127 | | | | | |
| | Prepost:LR | $\chi^2(1)$= 0.2 | 0.648 | | | | | |
| | Ablation:Prepost:LR | $\chi^2(2)$= 1.0 | 0.621 | | | | | |

Left (PA, n = 28), right (PA, n = 29), and both left and right ( PA, n = 13) sides.
Linear or generalized linear models were selected using Akaike's information criterion function to determine the best fit model for analyzing swimming distance, NOA, DIR, and DOR.
***: Significant at alpha = 0.001; **: significant at alpha = 0.01; *: significant at alpha = 0.05.
ANOVA was performed using the generalized linear models fitting function. Post-hoc tests were performed using the Wilcoxon signed-rank (paired) test followed by Holm's multiple-test correction.

### Supplementary Table 5. Statistical scores for Figure 4 and Figure 5—figure supplement 1.

| | **Generalized linear model** | | | **Wilcoxon signed-rank (paired)** | | | **Holm's correction** | |
|---|---|---|---|---|---|---|---|---|
| | | Statistics | p-value | | statistics | p-value | alpha = 0.05 | |
| *Swimming Distance* | Population (SF, TI, PA) | F(2, 106) = 67.214 | < 2E-16 | Before vs. After (SF) | V = 37 | 0.117 | 0.233 | K= 3 |
| | Starvation (Before, After) | F(1, 106) = 7.417 | 7.56E-03 | Before vs. After (TI) | V = 207 | 9.54E-06 | **2.86E-05*** ** | |
| | Pop:Starvation | F(2, 106) = 10.199 | 8.89E-05 | Before vs. After (PA) | V = 106 | 0.985 | 0.985 | |
| *Number of Approaches (NOA)* | LR (Left, Right) | F(1, 212) = 0.033 | 0.855 | Before vs. After (SF) | V = 8.5 | 3.74E-03 | **0.011*** | K= 3 |
| | Population (SF, TI, PA) | F(2, 212) = 102.928 | <2E-16 | Before vs. After (TI) | V = 81 | 0.846 | 0.846 | |
| | Starvation (Before, After) | F(1, 212) = 0.000 | 0.989 | Before vs. After (PA) | V = 134 | 0.286 | 0.573 | |
| | LR:Pop | F(2, 212) = 0.128 | 0.880 | | | | | |
| | LR:Starvation | F(1, 212) = 0.144 | 0.705 | | | | | |
| | Pop:Starvation | F(2, 212) = 4.485 | 0.012 | | | | | |
| | LR:Pop:Starvation | F(2, 212) = 0.050 | 0.951 | | | | | |
| *Duration Spent Inside the Radius (DIR)* | LR (Left, Right) | F(1, 212) = 9.469 | 2.37E-03 | Before vs. After (SF) | V = 7 | 1.76E-03 | **5.27E-03**** | K= 3 |
| | Population (SF, TI, PA) | F(2, 212) = 71.014 | <2E-16 | Before vs. After (TI) | V = 183 | 2.33E-03 | **4.65E-03**** | |
| | Starvation (Before, After) | F(1, 212) = 2.856 | 0.092 | Before vs. After (PA) | V = 87 | 0.522 | 0.522 | |
| | LR:Pop | F(2, 212) = 0.415 | 0.661 | L vs. R (SF, Before) | V = 39 | 0.144 | 0.144 | K= 6 |
| | LR:Starvation | F(1, 212) = 0.122 | 0.728 | L vs. R (SF, After) | V = 20 | 0.025 | 0.099 | |
| | Pop:Starvation | F(2, 212) = 8.340 | 3.26E-04 | L vs. R (TI, Before) | V = 11 | 1.05E-04 | **6.29E-04*** ** | |
| | LR:Pop:Starvation | F(2, 212) = 0.002 | 0.998 | L vs. R (TI, After) | V = 13 | 2.86E-03 | **0.014*** | |
| | | | | L vs. R (PA, Before) | V = 48 | 0.035 | 0.105 | |
| | | | | L vs. R (PA, After) | V = 52 | 0.048 | 0.097 | |
| *Duration Spent Outside the Radius (DOR)* | LR (Left, Right) | F(1, 212) = 7.281 | 7.53E-03 | Before vs. After (SF) | V = 131 | 3.05E-04 | **9.16E-04*** ** | K= 3 |
| | Population (SF, TI, PA) | F(2, 212) = 5.405 | 5.14E-03 | Before vs. After (TI) | V = 69 | 0.185 | 0.370 | |
| | Starvation (Before, After) | F(1, 212) = 0.379 | 0.539 | Before vs. After (PA) | V = 124 | 0.498 | 0.498 | |
| | LR:Pop | F(2, 212) = 5.269 | 5.84E-03 | L vs. R (SF, Before) | V = 63 | 0.821 | | |
| | LR:Starvation | F(1, 212) = 0.264 | 0.608 | L vs. R (SF, After) | V = 82 | 0.495 | | |
| | Pop:Starvation | F(2, 212) = 0.634 | 0.531 | L vs. R (TI, Before) | V = 65 | 0.143 | | |
| | LR:Pop:Starvation | F(2, 212) = 0.237 | 0.789 | L vs. R (TI, After) | V = 71 | 0.216 | | |
| | | | | L vs. R (PA, Before) | V = 84 | 0.452 | | |
| | | | | L vs. R (PA, After) | V = 95 | 0.729 | | |

Surface fish (SF, n = 16),Tinaja cavefish (TI, n = 20), and Pachón cavefish (PA, n = 20).
Linear or generalized linear models were selected using Akaike's information criterion function to determine the best fit model for analyzing swimming distance, NOA, DIR, and DOR.
***: Significant at alpha = 0.001; **: significant at alpha = 0.01; *: significant at alpha = 0.05.
ANOVA was performed using the generalized linear model fitting function. Post-hoc tests were performed using the Wilcoxon signed-rank (paired) test followed by Holm's multiple-test correction.

#### Supplementary Table 6. Statistical scores for Figure 5C and D.

| | General Mixed-Effects Mode (Gamma, Random effect = individual fish) | | | General Mixed-Effects Mode (Gamma,Random effect = individual fish) | | | Holms correction | |
|---|---|---|---|---|---|---|---|---|
| | | Statistics | | | statistics | p-value | alfa = 0.05 | |
| *Bias Index (NOA)* | Population (SF, TI, PA) | $X^2(2)$= 1.667 | 0.436 | Before vs After (SF) | $X^2(1)$= 0.011 | 0.917 | | K= 3 |
| | Starvation (Before, After) | $X^2(1)$= 0.300 | 0.584 | Before vs After (TI) | $X^2(1)$= 10.278 | 1.35E-03 | **4.04E-03**** | |
| | Pop:Starvation | $X^2(2)$= 7.532 | 0.023 | Before vs After (PA) | $X^2(1)$= 0.267 | 0.605 | | |
| *Bias Index (NOA)* | Population (SF, TI, PA) | $X^2(2)$= 10.945 | 0.042 | Before vs After (SF) | $X^2(1)$= 0.006 | 0.936 | | K= 3 |
| | Starvation (Before, After) | $X^2(1)$= 0.540 | 0.462 | Before vs After (TI) | $X^2(1)$= 5.245 | 0.022 | 0.066 | |
| | Pop:Starvation | $X^2(2)$= 5.182 | 0.075 | Before vs After (PA) | $X^2(1)$= 0.198 | 0.656 | | |

Surface fish (SF, n= 17),Tinaja (TI, n= 9), and Pachón (PA, n= 20).
Linear or generalized linear models were selected using the Akaike's information criterion function to address the best fit model to analyze bias index for NOA and DIR.
***: Significant at alpha = 0.001
**: Significant at alpha = 0.01
*: Significant at alpha = 0.05

Analysis of variance was done with the General Mixed-Effects Model fitting function. Post-hoc tests were performed using the General Mixed-Effects Model by Holm's multiple-test correction.

#### Supplementary Table 7. Correlation analysis for Figure 4.

| | **SURFACE FISH** | | **TINAJA** | | **PACHÓN** | |
|---|---|---|---|---|---|---|
| | **Kendall Tau** | **P-value** | **Kendall Tau** | **P-value** | **Kendall Tau** | **P-value** |
| **Correlation against total NOA (Before)** | | | | | | |
| Left SN number | 0.226 | 0.236 | 0.053 | 0.762 | 0.226 | 0.182 |
| Right SN number | 0.175 | 0.361 | -0.100 | 0.568 | 0.173 | 0.323 |
| **Correlation against total NOA (After)** | | | | | | |
| Left SN number | −0.379 | 0.052 | 0.267 | 0.124 | −0.030 | 0.861 |
| Right SN number | **−3.292** | **9.96E-04***** | 0.104 | 0.548 | −0.113 | 0.518 |
| **Correlation against increased amounts of total NOA (After - Before)** | | | | | | |
| Left SN number | −0.315 | 0.094 | 0.240 | 0.163 | **−0.394** | **0.020*** |
| Right SN number | −0.352 | 0.063 | 0.182 | 0.288 | −0.340 | 0.052 |

***: Significant at alpha = 0.001
**: Significant at alpha = 0.01
*: Significant at alpha = 0.05
